# Chemically Induced Nuclear Pore Complex Protein Degradation via E3 Ligase TRIM21

**DOI:** 10.1101/2024.12.03.626577

**Authors:** Xiaomei Li, Qingyang Wang, Anping Guo, Yaping Qiu, Qiuxia Chen, You Li, Lanjun Zhang, Yaxin Guo, Xiaoyun Meng, Shiqian Li, Guizhi Liu, Liyun Zhang, Jian Liu, Xianyang Li, Longying Cai, Xuemin Cheng, Chuan Liu, Xiaotao Wang, Andrew Wood, James Murray, Guansai Liu, Jin Li, Xiaodong Huang, Dengfeng Dou

## Abstract

Despite the exciting progress of the bifunctional degrader molecules, also known as proteolysis-targeting chimeras (PROTACs), the rapidly expanding field is still significantly hampered by the lacking of available E3 ligase ligands. Our research bridges this gap by uncovering a series of small-molecule ligands to the E3 ligase TRIM21 through DNA-Encoded Library (DEL) technology. We confirmed their interaction with TRIM21 using crystallography and demonstrated their anti-proliferative effects across various cancer cell types. Furthermore, proteomic studies identified that the mRNA Export Factor GLE1, and the Nuclear Pore Complex Protein NUP155, were significantly down-regulated upon TRIM21 ligand treatment. This degradation required TRIM21 and was ubiquitin-proteasome-dependent. More specifically, NUP155 was the primary target for the TRIM21 ligands, while GLE1 was considered a passenger target upon the initial degradation of NUP155. Using immunofluorescence techniques, we further demonstrated that the degradation of GLE1 and NUP155 proteins impaired the integrity of the nuclear envelope, leading to cell death. Highlighted by this research, a novel mode of action has been discovered for the TRIM21 E3 ligase ligand, acting as a monovalent degrader that triggers de novo interaction with functional complex proteins and induces their degradation.

## Introduction

Governed by the ubiquitin-proteasome system (UPS), E3 ubiquitin ligases represent a highly targeted machinery of protein post-translational modification, which specifically and catalytically transfers ubiquitin molecules onto substrate lysine residues, and leads to subsequent proteasomal degradation of their substrates^1^. E3 ligases mutations may negatively affect their activities, for instance the loss-of-function mutation of the F-box and WD repeat domain containing 7 (FBXW7)^2^, or overexpression that promotes the degradation of their substrates including tumor suppressors, both of which are found to be tightly associated with oncogenic transformation and cancer progression^3,4^. In the latter case, development of molecules that inhibit those tumor-specific overexpressing E3 ligases, such as the mouse double minute 2 homologue (MDM2)^5^ and inhibitors of apoptosis (IAPs)^6^, could contribute to cell proliferation inhibition and represent potential therapeutic solutions. Alternatively, hijacking and redirecting E3 ligases to neo-substrates and selectively remove disease-causing proteins represents another attractive therapeutic strategy. Such induced targeting protein degradation (TPD) strategies enabled by mono-functional Molecular Glues (MGs) or bifunctional Proteolysis Targeting Chimeras (PROTACs) provide an additional layer of regulatory specificity, which can be harnessed to modulate the UPS for therapeutic purposes^7–9^.

Although the human genome encodes over 600 E3 ubiquitin ligases^10^, most of these enzymes are yet to be fully characterized and only very few reliable ligands are available to re-purpose these ligases, limiting the scope of chemically induced TPD^11^. The two most commonly explored E3 ligases are Cereblon (CRL4^CRBN^) and von Hippel-Lindau (CRL2^VHL^), both of which belong to the multi-subunit Cullin-RING E3 ubiquitin ligase (CRL) family^12,13^. Furthermore, the design of PROTAC molecules is still largely empirical and optimization is required to pair a right E3 ligase with an intended target protein, warranting further exploration of potential ligands for more E3 ligases. The tripartite motif-containing (TRIM) family of E3 ligases, characterized by their unique tripartite motif, represent one of the largest RING type ligases^14^. Unlike the multi-subunit CRL family, the TRIM family ligases contain both substrate recognition and ubiquitination catalytic domain in a single protein. Despite the compelling evidences on their cellular functions at multiple regulatory levels^15,16^, efforts on TRIM E3 ligase ligand exploration has been minimal. One such study has reported the identification of a small molecule ligand to the PRY-SPRY domain of TRIM58 with micromolar affinity^17^. Other studies have targeted a bromo-domain containing TRIM24^18,19^, with the optimized ligand achieving sub-micromolar affinity. Among all TRIM family members, TRIM21 is of particular interest as it behaves as an intracellular antibody receptor by interacting with the Fc domain of antibodies^20,21^. TRIM21 harbors a RING, B-Box, coiled-coil, and PRY-SPRY domains, with its RING domain catalyzes the transfer of ubiquitin, while the PRY-SPRY domain is used for substrate recognition. Each TRIM21 monomer undergoes dimerization through its coiled-coil domain to be active for cellular functions. As an antibody receptor, TRIM21 was leveraged for protein degradation in a ‘TRIM-Away’ approach, where the substrate protein was recognized via its antibody and then recruited to TRIM21^22,23^. Furthermore, neo-substrate degradation was achieved through an engineered biomolecule that incorporated a target protein binding antibody to the N-terminal Ring-B-box-coiled-coil (RBCC domain) of TRIM21^25^. All these studies suggest that TRIM21 is a highly processive E3 ligase that could be potentially redirected to a neo-substrate by PROTAC molecules and promote the degradation of target proteins of interest. Therefore, identification of ligands to TRIM21 could not only offer insights into the complex mechanisms of the UPS, but also potentially pave a way for novel TPD-driven therapeutic interventions.

Here we utilized DNA Encoded Library (DEL) screening technology^24^ to identify a series of TRIM21 ligands, specifically the ligands for the TRIM21 PRY-SPRY domain, the substrate interaction domain. Surprisingly, the identified monovalent TRIM21 ligands could inhibit the growth of a set of cancer cells through degradation of nuclear pore proteins and disruption of the nuclear pore integrity. Therefore, the TRIM21 ligand unveiled a novel mode of action in which it acts as a monovalent degrader, capable of redirecting TRIM21 E3 ligase activity towards the degradation of neo-substrates.

## Results and Discussion

### TRIM21 Ligand Identification and Binding Confirmation

Both His- and Strep II-tagged TRIM21^PRY-SPRY^ (287-465aa) proteins were used in DEL affinity selection. The two protein constructs afforded strong signals with a high degree of overlap, as demonstrated by a correlation analysis yielding a correlation coefficient (r) of 0.946 (Figure S. 1a). The detailed selection result was presented in DataWarrior cubes for individual DEL, with each axis representing the building blocks used in each DEL synthesis cycle (Figure S. 1b,c). A representative cubic view highlighted the enrichment of DEL compounds exhibiting high sequence counts was shown in Figure. 1a. The line features with variation on the bubble size indicated the potential different contribution of building blocks on the binding signal. Compound **1** (Figure. 1b) was synthesized and validated with a dissociation constant (*K*_D_) of 193 nM by Surface Plasmon Resonance (SPR) towards Strep II-tagged TRIM21 proteins (Figure. 1c).

**Figure 1.**
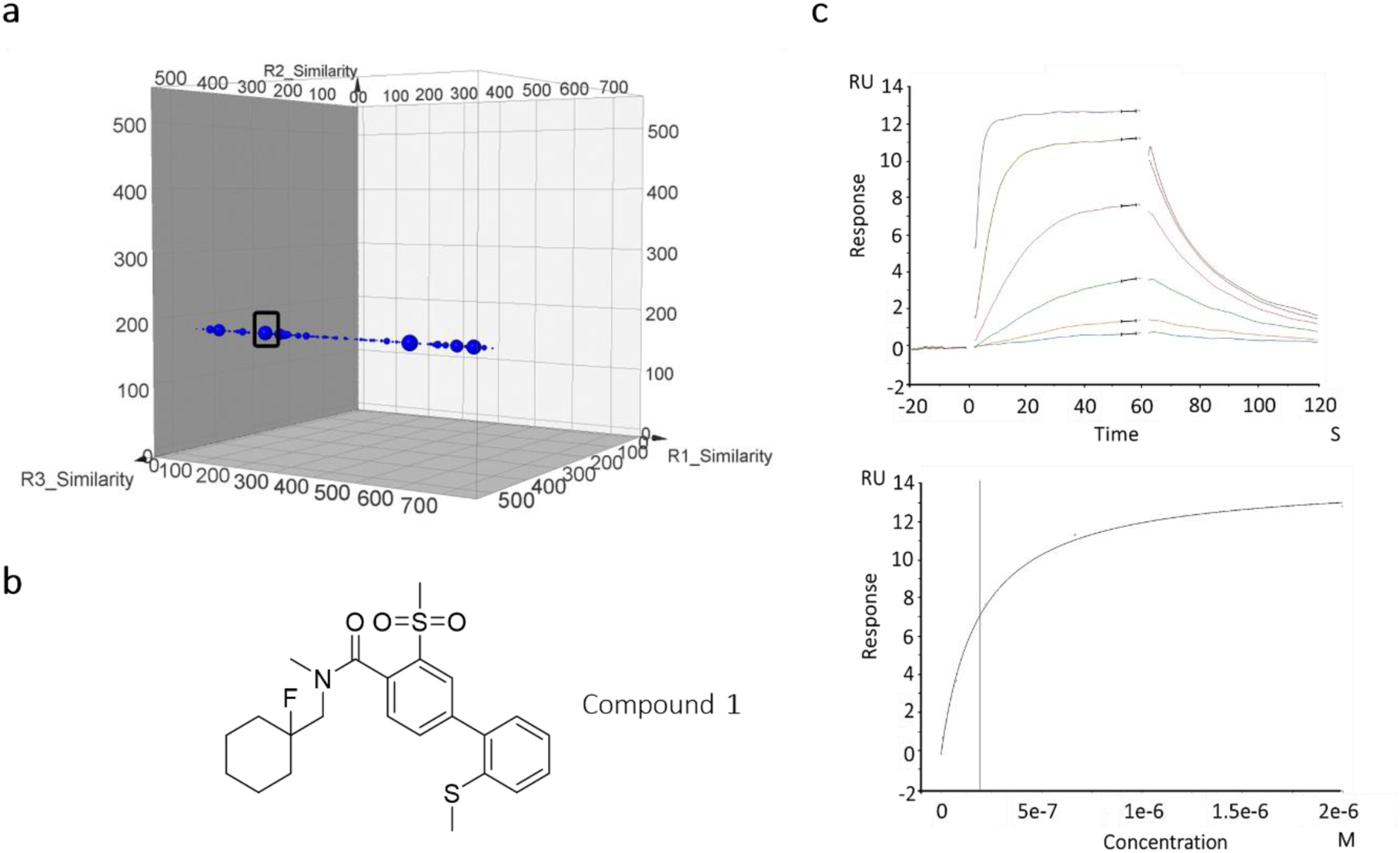
TRIM21 ligand identification from DEL selection and binding confirmation **a**, Selected line feature and compound enriched for TRIM21 DEL selection. **b**, Structure of DEL initial hit - Compound **1**. **c**, Confirmation of Compound **1** binding with Strep II-tagged TRIM 21 using Surface Plasmon Resonance (SPR)

### Analysis of Co-crystal Structure of TRIM21^PRYSPRY^ in Complex with Compound 1

To gain structural insight of TRIM21 ligand binding pose, we obtained co-crystal structure of the truncated TRIM21^PRY-SPRY^ (287-465aa) in complex with Compound **1** at 1.50Å resolution (Table 1). Specifically, Compound **1** binds snugly within a narrow, elongated pocket, fitting well with the shape of the pocket (Figure. 2a). The 2-methylthiophenyl moiety of Compound **1** extended into a hydrophobic sub-pocket formed by Y328, Fd450, D355, W381 and W383 with 109.4°bent in methylthio group (Figure. 2b). In addition, the methylsulfonyl benzene core forms π−π interactions with W381. No significant interaction was detected for the two oxygen atoms from the methylsulfonyl but the shape of the C-S-C bond was with a 100.8°angle. Two scaffold phenyl groups were twisted at a dihedral angle of 55.5 °to optimize their confirmation alignment within the pocket. Extended polar interaction (H-bond) was formed between the amide bond carbonyl with the backbone NH of L371. The tail cyclohexane occupying within the hydrophobic pocket by the two loop strands. In general, Compound **1** fits well into the topology of the TRIM21 hydrophobic pocket, enhancing binding through a series of polar and nonpolar interactions.

**Figure 2.**
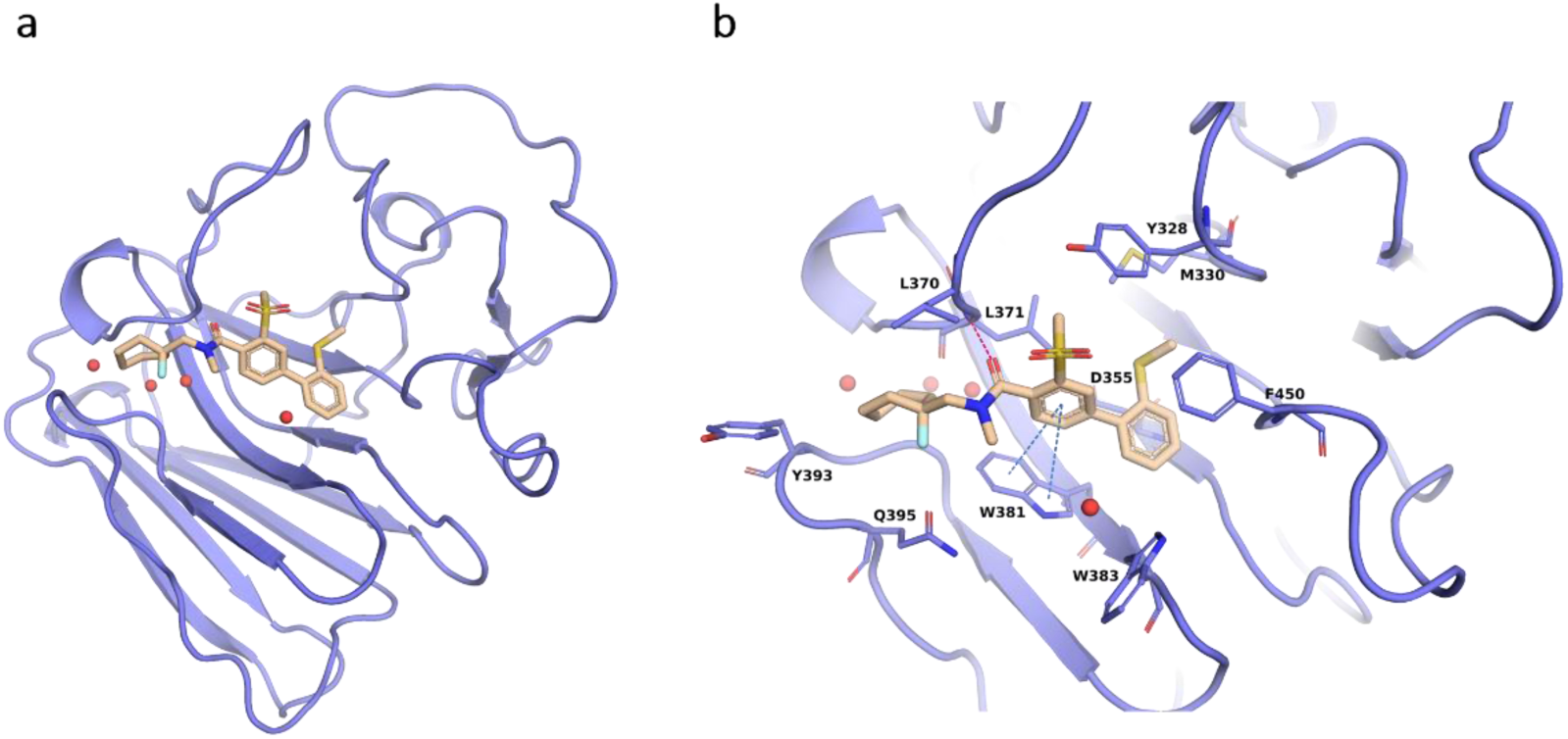
Co-crystal structure of TRIM21^PRYSPRY^ domain in complex with Compound **1**. a) X-ray co-crystal structure of TRIM21^PRYSPRY^ (light purple) with compound **1** (wheat sticks), **PDB ID: 9II5**. **b**, Compound **1** engages via apolar π−π interactions (marine dotted line) and polar interactions (red dotted lines) with several TRIM21 residues.

**Table 1.**
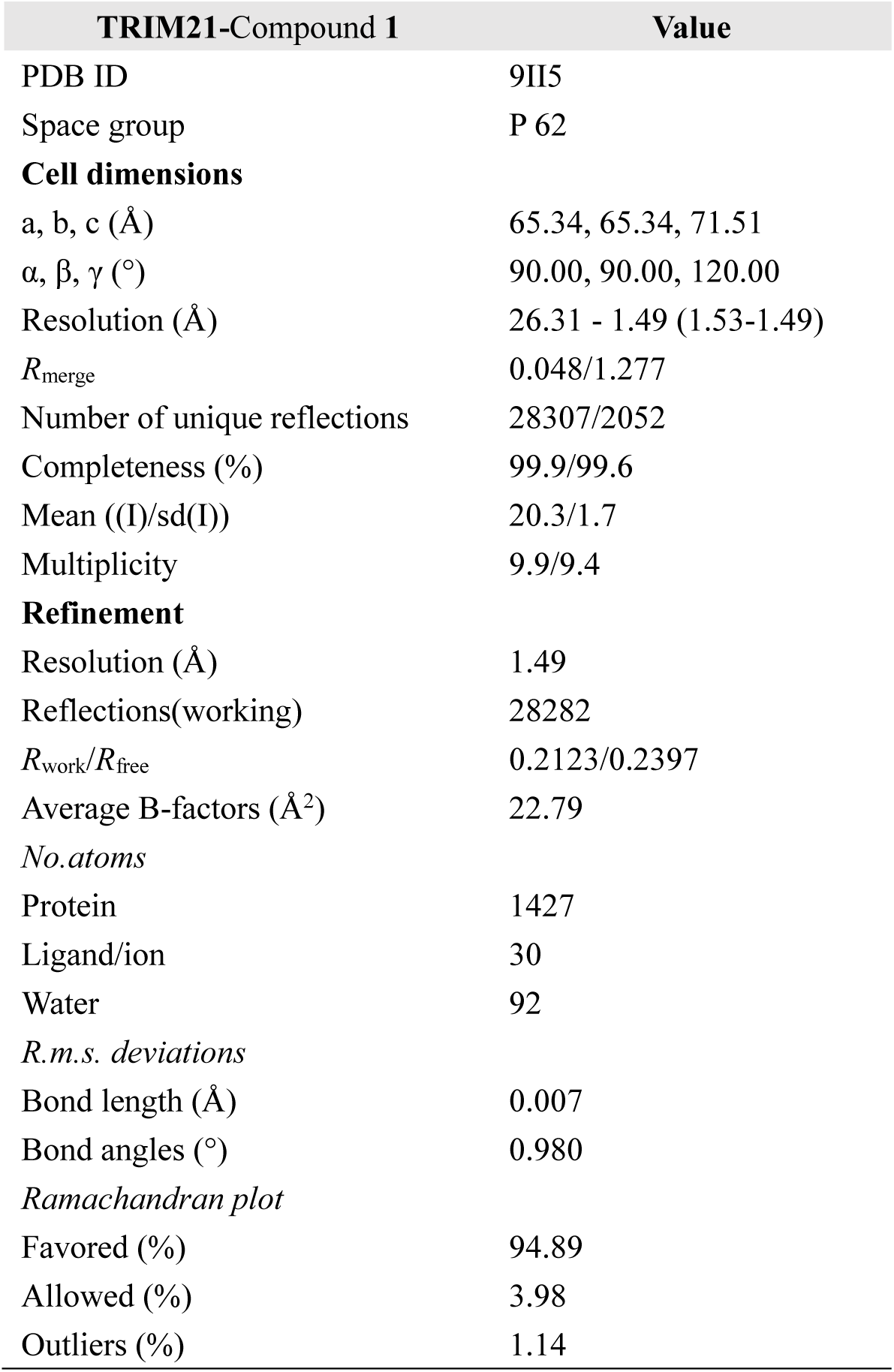
Data collection and refinement statistics.

| TRIM21-Compound 1 | Value |
| --- | --- |
| PDB ID | 9II5 |
| Space group | P 62 |
| **Cell dimensions** |  |
| a, b, c (Å) | 65.34, 65.34, 71.51 |
| $\alpha$, $\beta$, $\gamma$ (°) | 90.00, 90.00, 120.00 |
| Resolution (Å) | 26.31 - 1.49 (1.53-1.49) |
| $R_{\text{merge}}$ | 0.048/1.277 |
| Number of unique reflections | 28307/2052 |
| Completeness (%) | 99.9/99.6 |
| Mean ((I)/sd(I)) | 20.3/1.7 |
| Multiplicity | 9.9/9.4 |
| **Refinement** |  |
| Resolution (Å) | 1.49 |
| Reflections(working) | 28282 |
| $R_{\text{work}}/R_{\text{free}}$ | 0.2123/0.2397 |
| Average B-factors (Å<sup>2</sup>) | 22.79 |
| *No.atoms* |  |
| Protein | 1427 |
| Ligand/ion | 30 |
| Water | 92 |
| *R.m.s. deviations* |  |
| Bond length (Å) | 0.007 |
| Bond angles (°) | 0.980 |
| *Ramachandran plot* |  |
| Favored (%) | 94.89 |
| Allowed (%) | 3.98 |
| Outliers (%) | 1.14 |

### TRIM21 Ligands Induced Cell Growth Inhibition

Based on DEL screening data analysis, a series of Compound **1** derivatives, Compound **1a-1g** were synthesized (Table 2; see Supporting Information for synthetic routes and Nuclear Magnetic Resonance analysis). The binding of Compound **1a-1g** was validated in SPR with *K*_D_ ranging from 0.011 to 0.581 µM. In an attempt to further evaluate the ligand function, a cellular anti-proliferation assay was conducted for Compound **1a-1g** in a pancreatic cell line PANC-1 (Figure. 3a,b). Both Compound **1** and **1a** could arrest the cell growth with half inhibitory concentration (IC_50_) at micromolar range (Table 2). The inhibition of growth cannot be fully explained by the ligand’s affinity for TRIM21, since compound **1b, 1c** and **1g** with more potent *K*_D_ do not necessarily result in increased growth inhibition to cells. In molecular structure perspective, the cyclohexyl group might contribute to the anti-proliferation effect since it was presented in both Compound **1** and **1a**. To further investigate the relationship between TRIM21 potency and cell growth inhibition, **R2** group was further optimized by the guidance of co-crystal structure and molecular docking (Table 3). Replacing the methylthio group with fluorine (**2b**) and methyl (**2c**) group led to a decrease of potency and cellular activity (Figure. 3c). Further optimization to compound **2e** yield favorable TRIM21 binding and cellular activity. We then selected Compound **2e** (hereafter referred to as **HGC652**) for further investigation.

**Figure 3.**
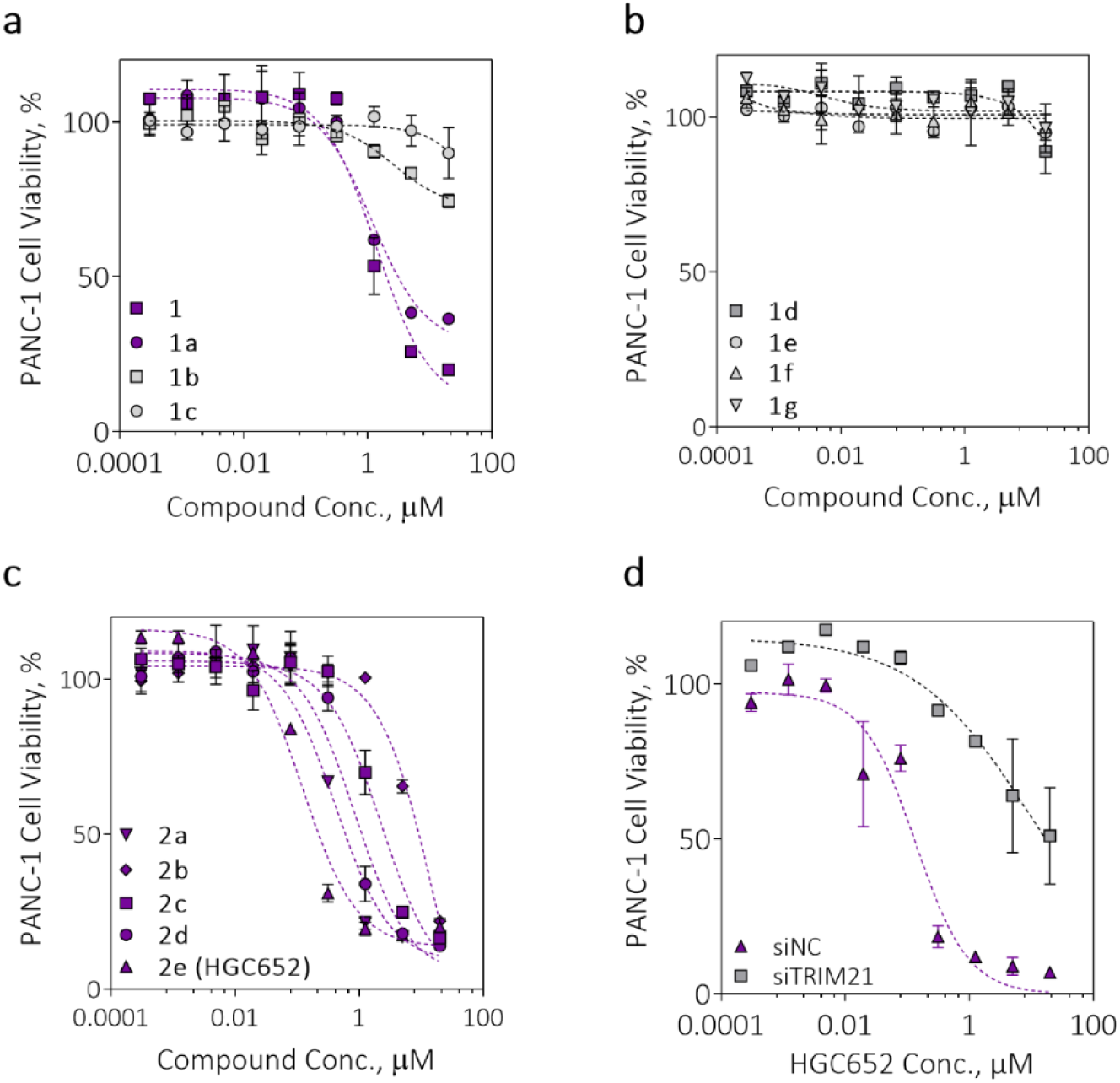
TRIM21 ligand affect cellular antiproliferation activity following 72 h of compound treatment (0∼20 µM). **a,** Growth inhibition effects of Compound **1, 1a-1c** on PANC-1 cells. **b,** Growth inhibition effects of Compound **1, 1a-1c** on PANC-1 cells. Compound **1** and **1a** exhibited inhibitory roles to the cell growth with IC_50_ values of 1.123 and 1.022 µM, respectively. **b,** Compound **1d-1g** did not induce growth inhibition effects to PANC-1 cells, with all IC_50_ > 20 µM. **c,** Compound **2a-2e** cause growth inhibition effects to PANC-1 cells, with IC_50_ values range 0.094 ∼ 5.138 µM. The most potent Compound **2e** was thereafter referred to **HGC652** in the following experiment. **d,** Growth inhibition of **HGC652** diminished in TRIM21 siRNA PANC-1 Cells. Cells were transiently transfected with TRIM21 siRNA for 48h following compound treatment for 72h.

**Table 2.**
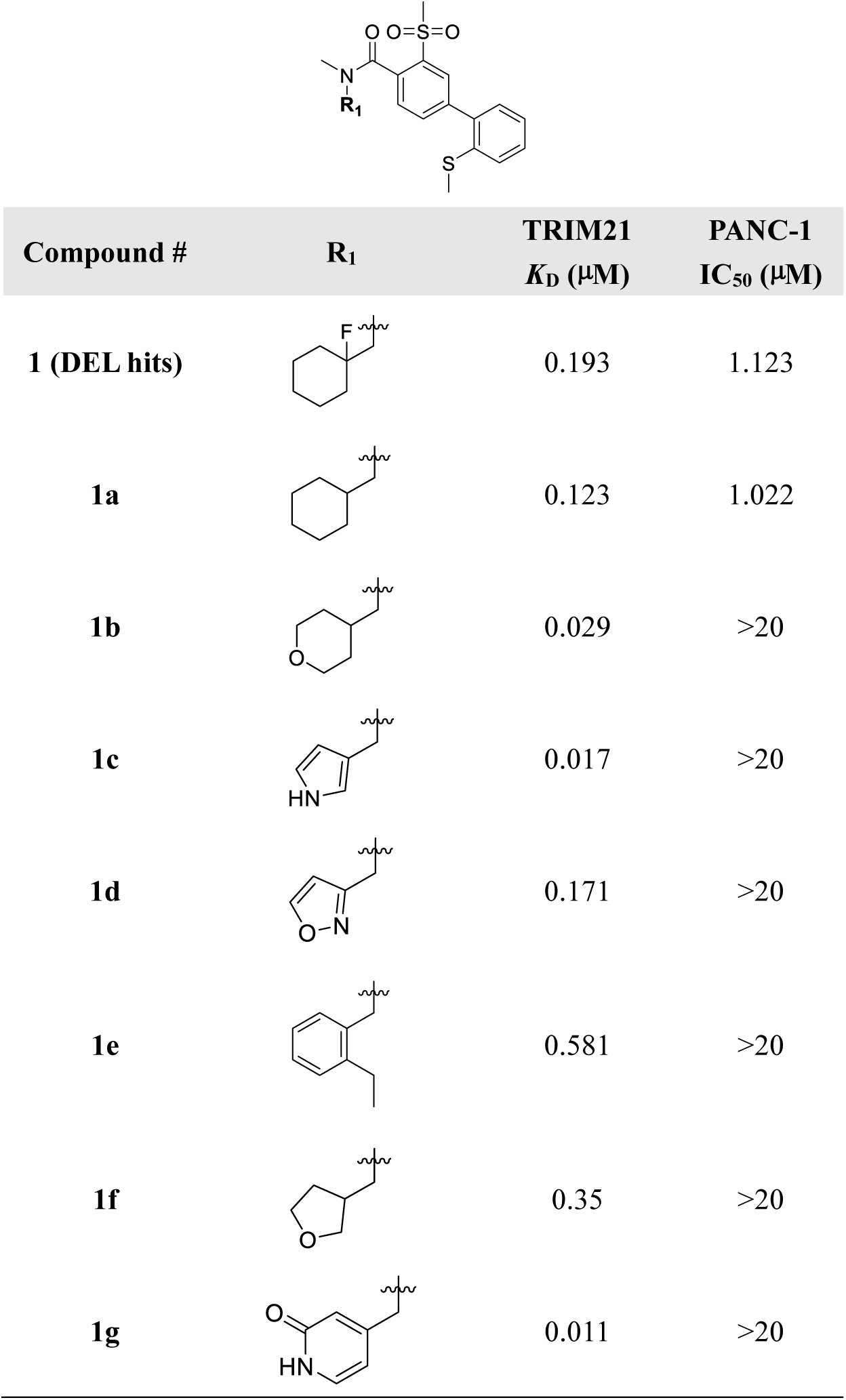
Structure –Activity Relationship of Compound 1. , **1a-1g**

| Compound # | R<sub>1</sub> | TRIM21<br>*K<sub>D</sub>* (μM) | PANC-1<br>IC<sub>50</sub> (μM) |
| --- | --- | --- | --- |
| **1** (DEL hits) |  | 0.193 | 1.123 |
| **1a** |  | 0.123 | 1.022 |
| **1b** |  | 0.029 | >20 |
| **1c** |  | 0.017 | >20 |
| **1d** |  | 0.171 | >20 |
| **1e** |  | 0.581 | >20 |
| **1f** |  | 0.35 | >20 |
| **1g** |  | 0.011 | >20 |

**Table 2.** Structure –Activity Relationship of Compound 2a-2e.

| Compound # | R<sub>1</sub> | R<sub>2</sub> | TRIM21<br>*K<sub>D</sub>* (μM) | PANC-IC<sub>50</sub> (μM) |
| --- | --- | --- | --- | --- |
| **2a** |  |  | 0.234 | 0.329 |
| **2b** |  |  | 2.15 | 5.138 |
| **2c** |  |  | 0.61 | 1.623 |
| **2d** |  |  | 1.57 | 0.791 |
| **2e (HGC652)** |  |  | 0.061 | 0.094 |

To confirm the engagement of TRIM21 in cell growth arrest, we first knocked down TRIM21 by siRNA in PANC-1 cells. The anti-proliferative effect of **HGC652** was significantly compromised following the knockdown of TRIM21 (Figure. 3d), suggesting that the inhibitory effect of **HGC652** is dependent on TRIM21 expression. Then the anti-proliferative effects of **HGC652** were further assessed using an array of tumor cell lines with varying levels of endogenous TRIM21 expression. Apart from the previously measured PANC-1 cells, eight other cancer cell lines demonstrated dose-dependent growth inhibition upon 72h compound treatment, with an IC_50_ range from 0.106 ∼ 0.822 μM (Figure. S2). In contrast, cells with low TRIM21 expression exhibited a limited response to the compound treatment. As anticipated, the varying cellular responses to **HGC652** treatment appeared to correlate with the level of endogenous TRIM21 expression. These findings suggest the potential for expanding the use of **HGC652** as an anti-cancer agent, specifically targeting cells with high TRIM21 expression.

### Proteomic and Transcriptomic Analysis Revealed a Subset of Nuclear Pore Complex Proteins as Potential Targets

TRIM21 has been shown to be a processive E3 ubiquitin ligase, we therefore conducted a time-resolved MS-based proteomics study, specifically targeting the proteomic alterations following **HGC652** treatment of PANC-1 cells. Adhering to a 1.5-fold change threshold^25^ and a p-value of less than 0.01, we uncovered 18 and 129 proteins exhibiting differential expression (hereafter referred to as DEPs), at 4- and 16-hours post-treatment, respectively (Figure. 4a,b). Notably, several nucleoporin (NUP) proteins featured prominently among the significantly down-regulated DEPs. To determine whether **HGC652** selectively targets nuclear pore-associated proteins for degradation, we utilized DeepLoc 2.0^26^ webserver to predict the subcellular localization of all DEPs from 16-hours post-treatment samples and a comparable number of non-DEPs, which was randomly chosen from proteins exhibiting no significant changes. The analysis indicated that the proportion of proteins primarily localized to the nucleus was approximately 12% higher among DEPs compared to the control protein set (Figure. 4c). Subsequently, we identified proteins classified as DEPs at both time intervals and examined their protein-protein interaction networks using the STRING database^27^, focusing exclusively on experimentally validated interactions with high confidence scores (greater than 0.7). Intriguingly, the analysis revealed that all differentially expressed NUPs engage in pairwise interactions, extending to interactions with GLE1 and AAAS (Figure. S3). To ascertain whether the observed downregulation of NUPs, GLE1, and AAAS was attributable to changes at the transcription levels, we conducted corresponding RNA-Seq experiments. These experiments confirmed no detectable decrease in mRNA transcripts (Figure. 4d), suggesting the decrease of these proteins was likely due to ubiquitination and degradation given that TRIM21 is a relatively very processive E3 ubiquitin ligase. Furthermore, Gene Ontology analysis of all DEPs from the 16-hour post-treatment samples suggested a significant enrichment in GO terms across biological processes, cellular components, and molecular functions, collectively suggesting that **HGC652** disrupts the integrity of a major nuclear-related complex (Figure. 4e).

**Figure 4.**
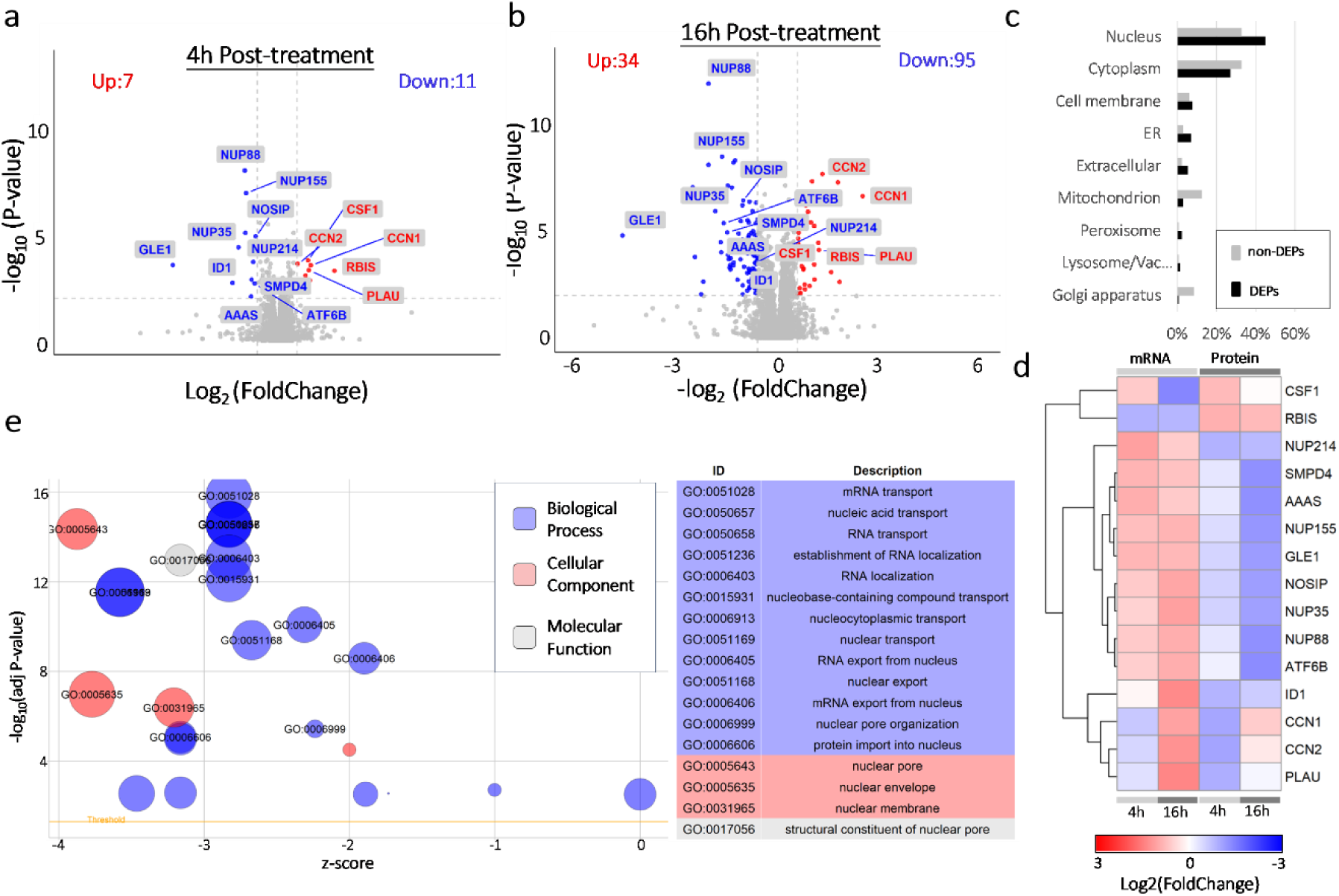
Proteomic and functional analysis of **HGC652** treatment. **a,** Quantitative proteomic data at 4-hour post-treatment in PANC-1 cells. **b,** Quantitative proteomic data at 4-hour post-treatment in PANC-1 cells. **c,** Proportions of proteins with varying primary subcellular localizations for DEPs versus selected control proteins from the 16-hour post-treatment samples. **d,** mRNA/protein expression levels of all DEPs common to both 4- and 16-hours post-treatment. **e,** Gene ontology analysis of DEPs from 16 hours post-treatment, high-lighting significant enrichment in biological processes, cellular components, and molecular functions.

### Validation of HGC652 Induced Nuclear Pore Complex Protein Degradation

We continued to investigate which key components of nuclear-related complex were degraded by **HGC652** treatment using immunoblotting assays. We started with a pair of interacting proteins GLE1 and NUP155^28^. Based on the proteomic data above, GLE1 protein showed the most significant time-dependent downregulation following compound incubation. NUP155 stands out due to its highest score within established protein-protein interaction networks (Figure. 4c). Indeed, after a 24-hour treatment, **HGC652** (0.5 and 5 μM) could induce a notable degradation of NUP155 and GLE1 proteins in PANC-1 cells (Figure. 5a). The application of proteasome inhibitor, MG132, but not autophagy-lysosome inhibitors Chloroquine and BafA1 (Figure. 5b), could effectively prevent NUP155 and GLE1 degradation, implying the engagement of a proteasome-dependent pathway. Furthermore, neither GLE1 (Figure. 5c) nor NUP155 (Figure. 5d) was degraded upon **HGC652** treatment when TRIM21 was knocked down, suggesting that TRIM21 is the bona fide E3 ubiquitin ligase responsible for the **HGC652**-mediated protein degradation in these cells.

**Figure 5.**
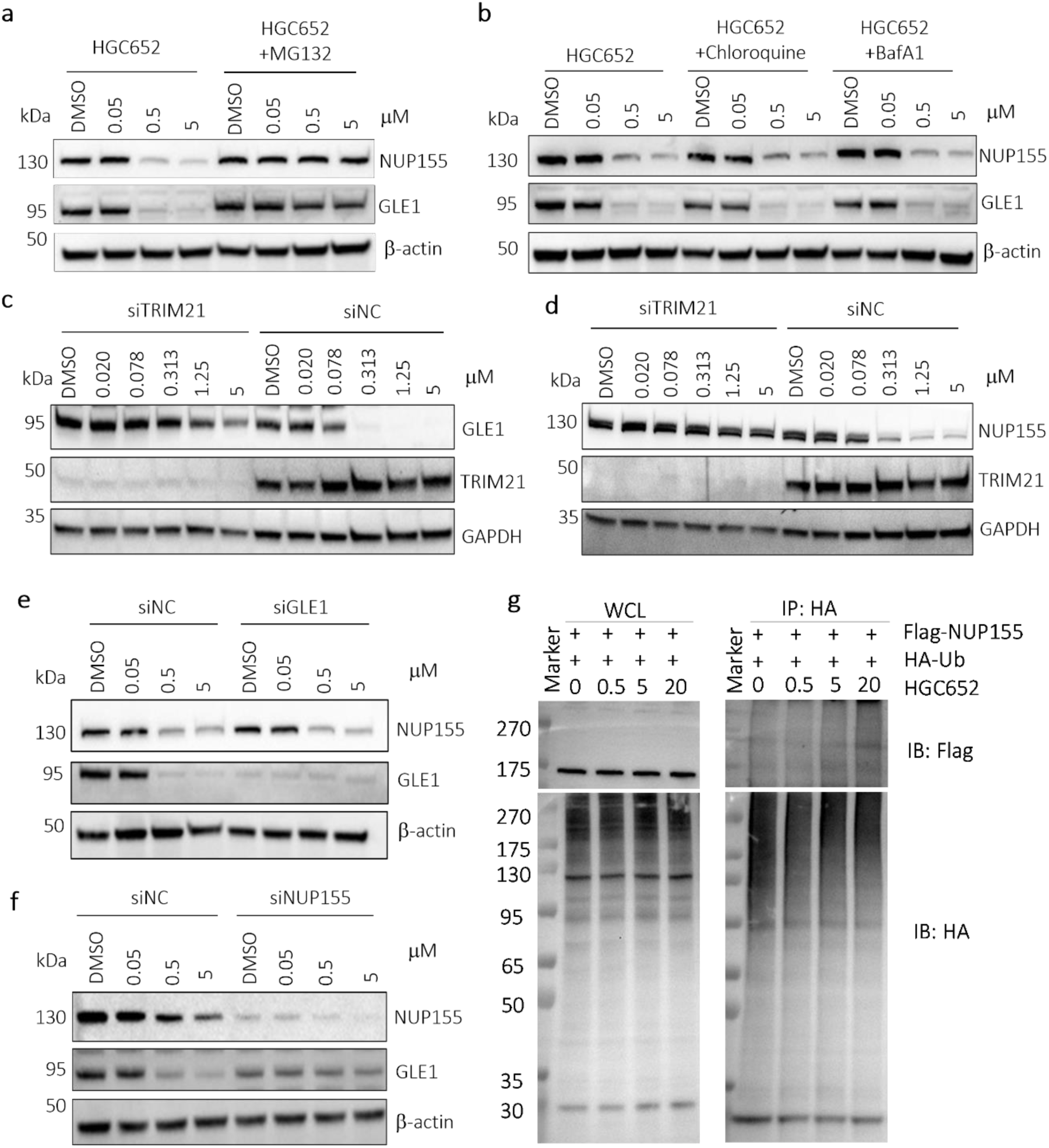
GLE1 and NUP155 protein degradation induced by TRIM21 ligand **HGC652**. **a,** Degradation of GLE1 and NUP155 at 24 hours of **HGC652** treatment in PANC-1 cells. Both GLE1 and NUP155 degradation was found to be dose-dependent, and could be rescued by proteasome inhibition. Cells were pre-treated with 5 μM MG132 for 0.5h, then co-treated with **HGC652** for another 16h. **b,** Neither GLE1 nor NUP155 degradation could be rescued by lysosome inhibition. Cells were pre-treated with 100 μM Chloroquine or 0.1 μM Bafilomycin A1 for 0.5h, then co-treated with **HGC652** for another 16h. **c,** Degradation of GLE1 at 24 hours of **HGC652** treatment in PANC-1 control and TRIM21 knock-down cells. **d,** Degradation of NUP155 at 24 hours of **HGC652** treatment in PANC-1 control and TRIM21 knock-down cells. **e, HGC652-**mediated NUP155 degradation was still observed in siGLE1 cells. **f, HGC652-**mediated GEL1 degradation was nearly abolished in siNUP155 cells. **g, HGC652** mediates polyubiquitination of NUP155 in HEK-293 cells. HEK-293 cells were transiently transfected with FLAG-NUP155 (3.2 μg) and HA-Ubiquitin (HA-Ub, 1.6μg) for 48h, and then pre-treated with MG132 (10 μM) for 0.5h. Then cells were treated with DMSO or HGC652 (0.5, 5, 20μM) in the presence of the proteasome inhibitor for another 2h before harvesting.

To identify which protein was degraded primarily by **HGC652**, we assessed NUP155 degradation where GLE1 was knocked down by siRNA. NUP155 degradation persisted even in the absence of GLE1 expression within the cells (Figure. 5e), suggesting that the **HGC652**-induced NUP155 degradation was independent of GLE1. On the contrary, the degradation of GLE1 was completely abrogated when NUP155 was absent (Figure. 5f). We further assessed NUP155 cellular ubiquitination level using HA-ubiquitin pull-down assays. Enhanced NUP155 ubiquitination was detected upon addition of **HGC652** (Figure. 5g), suggesting NUP155 degradation was triggered by UPS pathway.

During our ongoing research, a recent publication on TRIM21 suggests a similar finding on small molecule directed nuclear pore complex protein degradation via NUP98 autoproteolytic domain (APD)^29^. We hypothesized that our TRIM21 ligand **HGC652** may be conferred as the same mechanism, to first induce recruitment of NUP98 to TRIM21 and then trigger associated nuclear pore complex protein degradation (Figure. 6a). To verify such hypothesis, we employed FRET-based ternary complex assay using TRIM21^PRYSPRY^ and NUP98^APD^ proteins. With increasing concentrations of **HGC652**, a proximity induced FRET signals were detected between the two proteins (Figure. 6b), suggesting a molecular glue type of mode of action. We then analyzed the ternary complex formation using molecular modeling. Our data revealed that **HGC652** binds to the protein-protein interaction (PPI) interface between TRIM21^PRYSPRY^ and NUP98^APD^ (Figure. 6c,d). The interface exhibits favorable shape and electrostatic complementarity, and thus provides structural basis for a compound induced stabilizing complex. Previous studies have suggested that TRIM21 requires target-induced clustering for its E3 ligase activation^22^. The observed degradation in our study could be due to the natural assembly of the nuclear pore complex, providing structural basis for recruiting multiple TRIM21.

**Figure 6.**
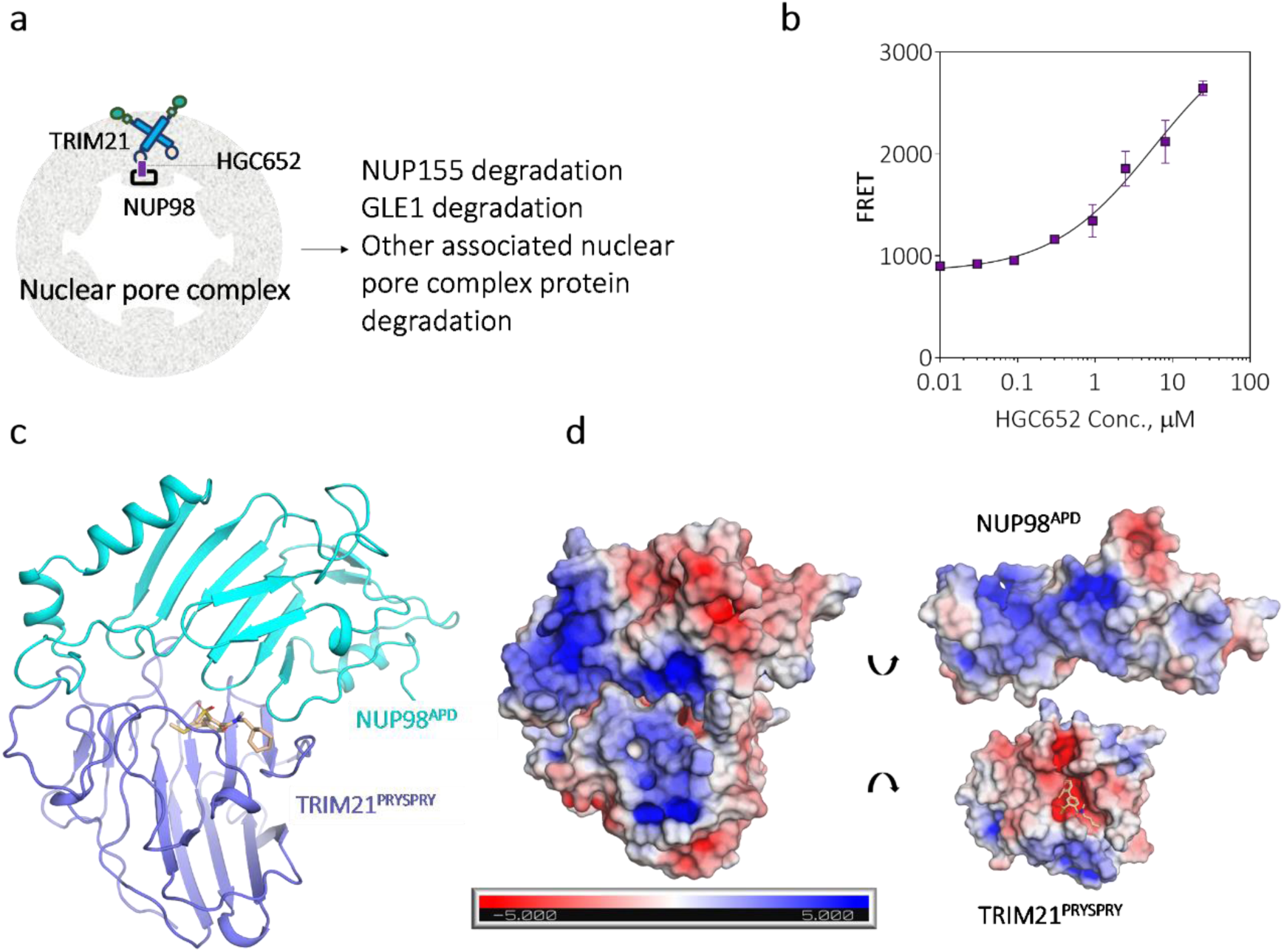
**HGC652** induced proximity between TRIM21 and NUP98. **a,** Hypothesis of **HGC652** mode of action. **b, HGC652** induced ternary complex formation between TRIM21^PRYSPRY^ and NUP98^APD^ proteins. **c, HGC652** and protein-protein docking studies on TRIM21 (**PDB ID: 9II5**) and NUP98 (**PDBID: 1KO6**, chain A). **d, HGC652** resides on Protein-protein interaction (PPI) interface between TRIM21 and NUP98 exhibiting favorable shape and electrostatic complementarity.

### Nuclear Envelope Impairment and Cell Death

Following a 24-hour incubation with **HGC652** in PANC-1 cells, there was a notable decrease in the number of foci detected by nuclear pore complex antibody mAb414 compared to the control cells (Figure. 7a). Alongside the reduction of the nuclear pore complex, PANC-1 cells showed changes in nuclear morphology and a decrease in DNA fluorescent signals, suggesting a possible genome instability due to the leakage of genetic material into the cytoplasm. We evaluated cell growth for 72h incubation period after transfection with NUP155, GLE1, and NC siRNA (48-h post-transfection), without any additional compound treatment. Cells transfected with NUP155 siRNA exhibited significant growth inhibition in comparison to the other two conditions (Figure. 7b), suggesting that NUP155 may have a more vital role in cell growth than GLE1. Considering the integral role GLE1 and NUP155 play in both the structural and functional aspects of the nuclear envelope^28,30^, our findings suggest that the TRIM21 ligand may disrupt these essential structures of the nuclear envelope, resulting in subsequent cell death, a similar effect that is consistent with previous studies involving NUP155 siRNA knock-down^31^.

**Figure 7.**
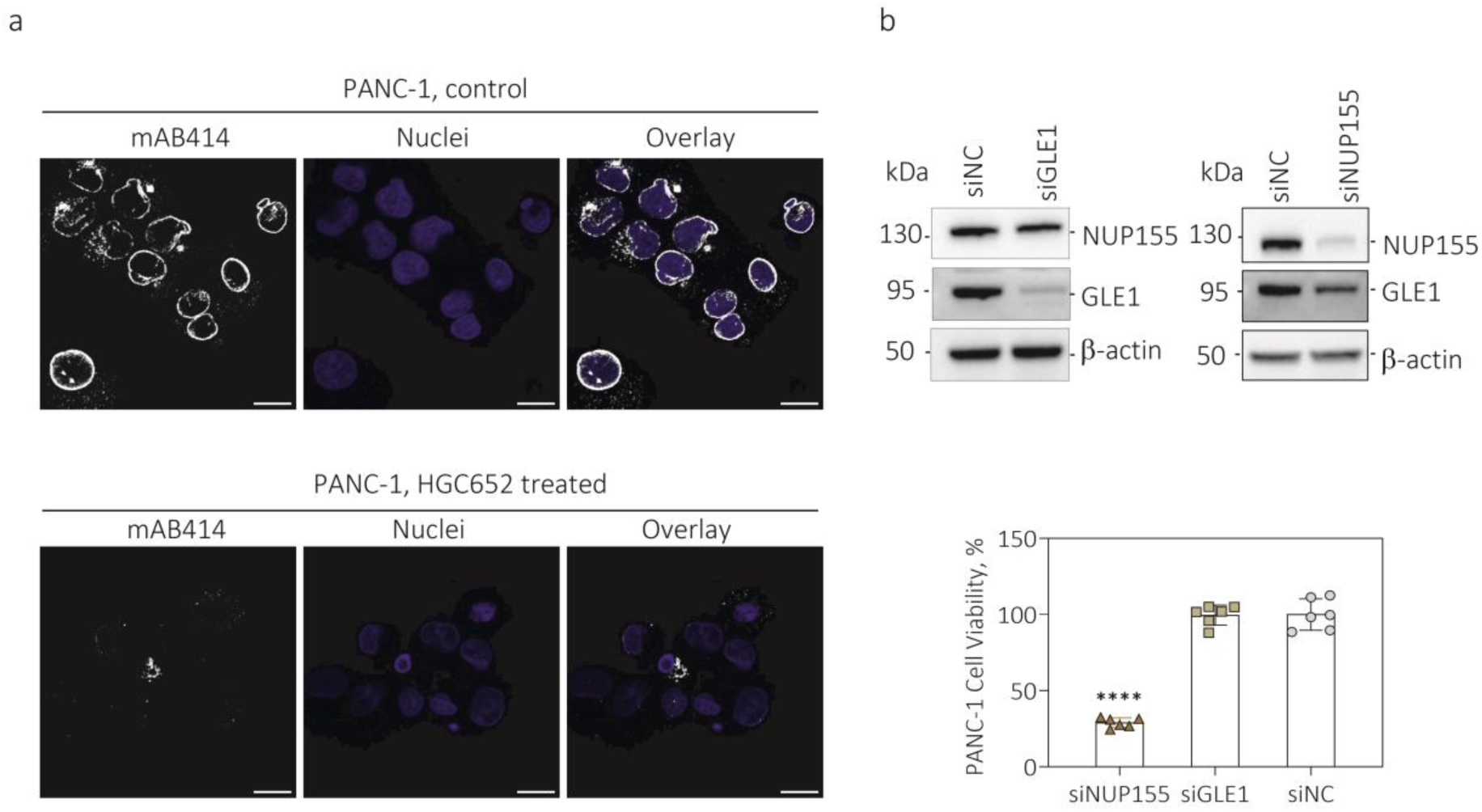
**HGC652** induced nuclear pore complex disintegration. **a,** mAB414 nuclear pore complex staining and nuclei staining in PANC-1 cells in the presence and absence of TRIM21 ligand **HGC652** treatment. Scale bar 20 µm. **b,** Cell viability of GLE1 and NUP155 siRNA knock-down (48h transfection) PANC-1 cells following a 72h culture period without additional compound treatment. Data were calculated as % of SiNC control samples. NUP155 knock-down showed significant reduction (\*\*\*\**P* < 0.0001) in comparison with siNC control group.

## Discussion

Our research has successfully led to discovery of novel monovalent molecules targeting E3 ligase TRIM21. The initial ligand, Compound **1** was identified via DEL screening, with *K*_D_ of 0.193 µM. Interestingly, a series of its close analogs exhibited TRIM21-dependent cellular effects to multiple cancer cells lines. Co-crystal analysis with Compound **1** allowed us to further optimize the ligands with improved TRIM21 binding and cellular activity. With the most potent compound, **HGC652** (*K*_D_ 0.061 µM, IC_50_ 0.094 µM), we unveiled that the ligand acts like a molecular glue molecule to induce proximity between TRIM21 and NUP98. Upon recruiting multiple TRIM21 onto nuclear pore complex, the degradation process potentially initiates with NUP155 and then GLE1, subsequently leading to the disintegration of the entire nuclear envelope. Such discovery of a chemically redirected E3 ligase activity towards noncanonical substrates, may provide novel insights into the UPS mechanism and stimulate further research in the field of targeted protein degradation.

## Materials and Methods

### DEL Selection

Both His-TRIM21^PRY-SPRY^ and Strep II-TRIM21^PRY-SPRY^ were used for selection against 455.51 billion encoded compounds (HitGen DEL), with the corresponding blank control included for comparison and background subtraction. Automated selection was carried out using a KingFisher Duo Prime Purification System (ThermoFisher) in a 96-well plate in a similar format as previously reported^32^. After equilibrated with selection buffer (7.6 mM NaH_2_PO_4_, 12.4 mM Na_2_HPO_4_, 150 mM NaCl, 0.3 mg/mL ssDNA, 0.1% Tween-20, pH 7.0, with 10 mM imidazole for Ni-charged MagBeads), Ni-charged MagBeads (GenScript, L00295) and Magstrep “type 3” XT Beads (IBA-Lifesciences, 2-4090-010) were transferred to a new row and incubated with 5 μM His-TRIM21^PRY-SPRY^ and Strep II-TRIM21^PRY-SPRY^ at room temperature (RT) for 30 min, respectively. Then, the immobilized protein along with the beads was incubated with DEL sample in selection buffer at RT for 1 h followed by 1 min wash at RT in 500 μL selection buffer (with 30 μM biotin for Magstrep “type 3” XT Beads groups) for five times. Retained DEL molecules were recovered by heat elution in elution buffer (7.6 mM NaH_2_PO_4_, 12.4 mM Na_2_HPO_4_, 300 mM NaCl, pH 7.0) at 95 °C for 10 min. The selection was repeated with the heat-eluted portion of the previous round and fresh protein for three times for His-TRIM21^PRY-SPRY^ and two times for Strep II-TRIM21^PRY-SPRY^, the recovered DEL samples were incubated with blank Ni-charged MagBeads and Magstrep “type 3” XT Beads respectively to further remove bead binders, then the output was amplified by PCR and sequenced by NovaSeq6000. Selection data were decoded, presented in DataWarrior (developed by Openmolecules) cubes, and analyzed by feature intensity using the PolyO score^33^.

### Surface Plasmon Resonance (SPR)

SPR measurement was carried out on a BIAcore T200 (Cytiva) at 25 °C in buffer containing 7.6 mM NaH_2_PO_4_, 12.4 mM Na_2_HPO_4_, 150 mM NaCl, 0.05% Tween-20, 1% DMSO, pH7.0. The Strep II-TRIM21^PRY-SPRY^ protein was captured on a S series CM5 sensor chip after the immobilization of Strep-Tactin XT by amine coupling. His-TRIM21^PRY-SPRY^ protein was captured on a S series NTA sensor chip. The binding affinity of the analyte was measured by passing serially increased concentrations of compound at the flow rate of 30 μL/min. The final response was calculated by double subtraction of the response from both reference channel and the zero concentration, binding affinity was fitted by steady state fitting model from the built-in software.

### Ternary Complex Assay

**HGC652** compound (0∼25 µM) was preincubated with 30 nM His-TRIM21^PRY-SPRY^ and 30 nM Strep II-NUP98^APD^ in assay buffer (20 mM PBS, 0.08% BSA, pH 7.0) at RT for 40 min. Then detection buffer (0.4nM Strep-tactin-Tb/ 13.34nM anti-6HIS-XL665) was added and incubated at RT for 1h. Fluorescence emission was measured at Ex=337nm and Em=620/665nm.

### Protein Expression

TRIM21^PRY-SPRY^ (residues 287-465) gene was cloned into pET28a vector for expression with a Thrombin cleavable His or twin strep tag at the N-terminus. For protein production, BL21 (DE3) cells were cultivated in TB medium supplemented with 50 μg/mL kanamycin at 37 °C until OD600 reached ∼1.4 and then cooled to 18 °C for 1 hour. Isopropyl 1-thio-D-galactopyranoside (IPTG) was added to 0.3 mM, and growth continued at 18 °C overnight. Cells were collected by centrifugation and pellets were suspended in Buffer A (20 mM Tris pH 8.0, 350 mM NaCl, 1 mM TCEP, 5% (V/V) glycerol, 0.05% CHAPs), lysed by high pressure homogenizer, 85 MPa, 3 times, and clarified by centrifugation at ∼15,000×g for 60 min.

The supernatant of His-TRIM21^PRY-SPRY^ and Strep II-TRIM21^PRY-SPRY^ was respectively loaded onto a 5 mL Nickel-Sepharose column and Strep-AC column, equilibrated in Buffer A. The column was washed with Buffer A after protein loading. The His-TRIM21^PRY-SPRY^ protein was eluted by an imidazole step gradient with Buffer A containing 10, 30, 50, 100, 300, 500 mM imidazole. The Strep II-TRIM21^PRY-SPRY^ was eluted with the Buffer A supplemented with 50 mM biotin (pH 8.0). The collected eluate was concentrated and further purified by size-exclusion chromatography using a HiLoad Superdex 200 16/600 gel filtration column in Buffer B (50 mM HEPES-HCl, 150 mM NaCl, 10%(v/v) glycerol, 1 mM TCEP, 0.05% w/v CHAPS, pH 8.0). Protein concentration was determined by the Bradford protein assay with bovine serum albumin (BSA) as a standard. The purified protein was stored in 50 mM HEPES (pH 8.0), 150 mM NaCl, 20% (V/V) Glycerol, 1 mM TCEP, 0.05% (W/V) CHAPs at -80 °C. For TRIM21^PRY-SPRY^ crystallization, tag cleavage was performed at 4 °C overnight, using the recombinant thrombin protease. The tag-free TRIM21^PRY-SPRY^ was further purified by the HiLoad Superdex 200 16/600 gel filtration column.

The coding regions of human NUP98 (729-880 aa) was synthesized and cloned into pET based vector with an N-terminal Strep II tag. Protein was overexpressed in BL21(DE3) cells induced with 0.3 mM IPTG. The tagged protein was captured by StrepTacin resin (IBA lifesciences) in a lysis buffer consisting of 50 mM Tris-HCl (pH7.5), 300 mM NaCl, 5% (v/v) Glycerol, 5 mM DTT and washed extensively with the lysis buffer. Protein was concentrated and subjected to a Superdex 200 increase 10/300GL gel filtration column (Cytiva lifesciences) in buffer containing 20 mM Tris-HCl, pH7.5, 300 mM NaCl, 10% (v/v) Glycerol, 5 mM DTT. NUP98 was concentrated to 1 mg/mL and stored at -80 °C.

### Determination of Co-crystal Structures of TRIM21^PRY-SPRY^/Compound 1 Complex

Compound **1** in 100% DMSO was added to TRIM21^PRY-SPRY^ protein solution at a 1:1.5 molar ratio and incubated on ice for approximately 4 hours. This mixture was centrifuged at 14,000 rpm for 10 minutes at 4 °C prior to crystallization. Crystallization was performed using sitting drop vapor diffusion. TRIM21^PRY-SPRY^ (at 8–10 mg/mL) was mixed with the reservoir solution at a 1: 1 volume ratio. TRIM21^PRY-SPRY^/Compound **1** cocrystals were observed in HR2-144 A7 containing 0.1 M citric acid (pH 3.5) and 2.8-3 M sodium chloride at 20°C after one day. Crystals were cryoprotected in the reservoir solution supplemented with 20% glycerol before flash-freezing in liquid nitrogen for data collection. These crystals were diffracted at 1.5 Å and belong to the space group P62 with unit cell parameters of a = 65.49, b =65.49, c= 71.75Å.

### Molecular Modeling

The analysis and plotting were done using PyMol. FRODOCK 3.0 was used for protein-protein docking with default parameters. The electrostatic potential molecular surface calculation was analyzed using APBS plugin of PyMol with default parameters.

### Cell Culture

A549, SK-N-AS, and U2OS cells were obtained from ATCC. A-431, PANC-1, MV-4-11, KURAMOCHI, MCF-7, HCC1806, MDA-MB-468 and NCI-H1373 cells were obtained from Cobioer. BxPC-3 cells were obtained from Jennio-bio. HEK-293 cells were obtained from Procell. MDA-MB-231 cells were obtained from National Collection of Authenticated Cell Cultures. A549, BxPC-3, KURAMOCHI, NCI-H1373, and HCC1806 cells were cultured in RPMI 1640 supplemented with 10% FBS and 100 μg/mL Normocin. MDA-MB-231 and U2OS cells were cultured in McCoy’s 5A supplemented with 10% FBS and 100 μg/mL Normocin. PANC-1 and MDA-MB-468 cells were cultured in DMEM supplemented with 10% FBS and 100 μg/mL Normocin. SK-N-AS cells were cultured in DMEM supplemented with 10% FBS and 1% NEAA. A431 cells were cultured in IMDM supplemented with 15% FBS and 1% penicillin-streptomycin solution. MV-4-11 cells were cultured in IMDM supplemented with 10% FBS and 100 μg/ml Normocin. HEK-293 cells were cultured in EMEM supplemented with 10% FBS and 100 μg/ml Normocin. All cells were maintained at 37 °C supplied with 5% CO_2_. Mycoplasma contamination was regularly examined using PCR Mycoplasma Detection Kit (G238, abm).

### Antibodies

The GLE1 antibody (PA5-117664), Alexa Fluor 488-conjugated goat anti-rabbit IgG (H+L) secondary antibody (A32731), Alexa Fluor 594-conjuated goat anti-mouse IgG (H+L) secondary antibody (A-11032) and Hoechst 33342 (H3570) were purchased from Invitrogen. NUP155 (66359-1-Ig) antibody and HRP-conjugated GAPDH (60004-1-Ig) were purchased from Proteintech. The HRP anti-beta-Actin (ab49900) and Anti-Nuclear Pore Complex mab414 (ab24609) antibodies were purchased from Abcam. The HRP-linked rabbit IgG (7074) and HRP-linked mouse IgG (7076) antibodies were purchased from Cell Signaling Technology. Anti-α-Tubulin antibody (T6199) was purchased from Sigma-Aldrich. Immobilon Western Chemiluminescent HRP Substrate (WBKLS0500) was purchased from Millipore. MG132 (S2619) was purchased from Selleck Chemicals.

### Cell Viability Assay

Cells were seeded in a 384-well clear bottom white plate (Corning) at a cell density of 750 cells per well and treated with compounds at the indicated concentrations for 3 days. Cell viability was assessed via the CellCounting-Lite 2.0 reagent (Vazyme). The luminescence signal was measured on a CLARIOstar (BMG LABTECH). The data were analyzed in GraphPad Prism 8.0.

### Functional analysis of Proteomics and RNA-Seq data

Proteomics data were generated with outsourcing facilities, and RNA sequencing experiments were performed in-house. The processing and quantification of RNA-Seq data were carried out utilizing a pipeline comprising HISAT2 for alignment, followed by StringTie for assembly and quantification of transcripts^34^. Differential expression analysis was conducted using the DESeq2 package^35^. The subcellular localization of selected proteins was predicted employing the DeepLoc-2.0 webserver, available at https://services.healthtech.dtu.dk/services/DeepLoc-2.0/. Analysis of protein-protein interactions was performed using the STRING database webserver (https://string-db.org/). The functional enrichment and Gene Ontology analysis of DEPs were conducted with the ClusterProfiler package^36^. Plots in this section were achieved using ggplot2 packages^37^ and the GOplot^38^, enabling the comprehensive illustration of the functional characteristics of the identified proteins.

### Cell Lysis and Immunoblotting Assays

Cells were collected and lysed in RIPA lysis buffer (Beyotime) supplemented with PMSF and protease inhibitor cocktail (Beyotime) on ice for 1 h. The supernatant was collected after centrifugation at 15,000 rpm for 10 min at 4 °C. Protein concentration was measured by BCA according to the manufacturer’s protocol (PA115, TIANGEN) and 4X LDS sample buffer was added.

Proteins were separated on 4-12% SDS-PAGE gels (GenScript) and transferred to PVDF membrane (0.2 μm, ISEQ00010, Millipore). Membrane was blocked with 5% milk in TBST for 1 hour at RT. The antibody was diluted with fresh 5% milk in TBST buffer (1:10,000 dilution for β-actin and GAPDH, 1:2,000 dilution for GLE1, NUP155) and incubated with membrane at 4 °C overnight. Membranes were washed three times with TBST buffer and incubated with secondary antibody (1:2,000 dilution in 5% milk in TBST) at room temperature for 1 h. After further washing, immunoblotting images were captured with an IBright-FL1500 image system.

### siRNA Transfection

The siRNAs targeting human TRIM21, NUP155and GLE1 were synthesized by GenePharma. Cells were seeded in 12-well plate and were transfected with siRNA for an indicated time using Lipofectamine RNAiMAX reagent (Invitrogen) according to the manufacturer’s protocol.

### In Cells Ubiquitination Assay

HA-ubiquitin and Flag-NUP155 expression plasmids were transfected with FuGENE® HD transfection reagent (E2311, Promega) in HEK293 cells. After 48h transfection, the cells were pre-treated with 10 μM proteasome inhibitor MG132 for 0.5h, then co-treated with HGC652 for another 2h before harvesting. The cells were lysed in RIPA lysis buffer supplemented with PMSF and protease inhibitor cocktail on ice for 0.5 h. Protein concentration was measured by BCA protein assay kit following the manufacturers’ protocol. Protein levels among samples were normalized to 2 μg/μL with ice-cold NP40 cell lysis buffer supplemented with PMSF and protease inhibitor cocktail. 475 μL of supernatant were mixed with 25 μL of 20% SDS, and heated at 95 °C for 10 min. The denature supernatant were diluted to 10 mL final volume with ice-cold NP40 cell lysis buffer supplemented with PMSF and protease inhibitor cocktail. 60 μL anti-HA magnetic bead (P2121, Beyotime) were added into supernatant and rotated overnight at 4 °C. The beads were washed five times with NP40 buffer. The washed anti-HA beads were resuspended in loading buffer and heated at 95 °C for 10 min. Protein samples were separated by SDS-PAGE, followed by immunoblot.

### Immunofluorescence

Cells were seeded and grown in 96 well glass bottom plate (P96-1.5H-N, Cellvis). After compound treatment, culture medium was discarded and replaced with 150 μl of 4% paraformaldehyde solution for 15 min at room temperature. The fixed cells were permeabilized with 150 μl 0.1%Triton X-100 solution for 15 min at room temperature, and then blocked with blocking reagent (Roche) for another 60 min at room temperature. The cells were incubated overnight at 4 °C with Nuclear Pore Complex mab414 antibody at 1:400 dilution in blocking reagent. Cells were washed three times with PBST buffer and incubated with Alexa Fluor 594-conjuated goat anti-mouse IgG (H+L) secondary antibody and Hoechst 33342 at 1:1,000 dilution at room temperature in dark for 2 h. Cells were washed three times with PBST buffer. The images were captured by a Zeiss LSM900.

## Author Contributions

XL, DD and XH conceived of the research and wrote the paper. QW and XW designed and synthesized TRIM21 compounds. AG, SL and GL performed cellular assays. YL for bioinformatic analysis. QC, JL, XC, CL for DNA-encoded library screening and data analysis. XL and LC characterized compounds in biophysical and biochemical assays. XM, LZ, YG expressed and purified proteins, and cocrystalization. YQ and LZ helped to analyze structural data. GL designed DNA-encoded library for screening. AW, JM and JL helped with the manuscript.

