## Supplemental data for "Chemically Induced Nuclear Pore Complex Protein Degradation via E3 Ligase TRIM21"

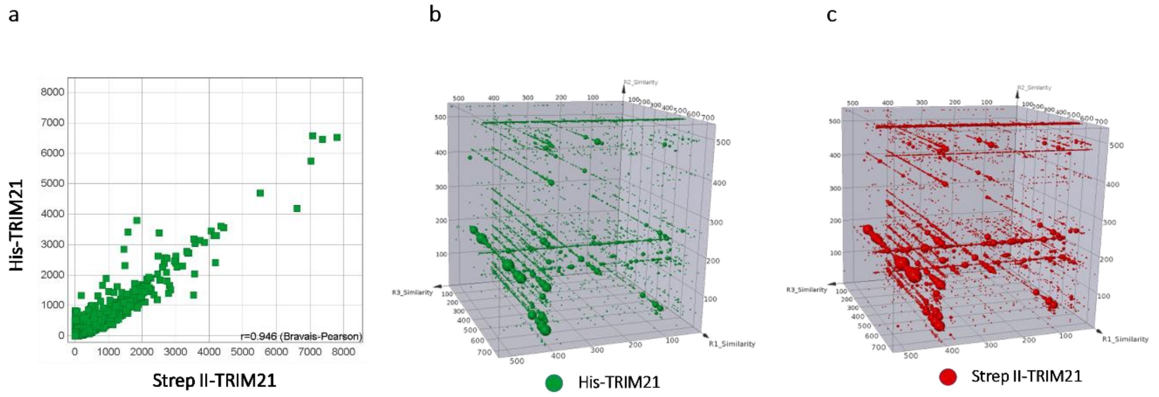


**Figure S1. TRIM21 DEL selection data.** **a**, Correlation analysis of DEL selection sequence count signals between His-tagged TRIM21 and Strep II-tagged TRIM21. **b**, Representative cubic view of His-tagged TRIM21 from DEL selection. **c,** Representative cubic view of Strep II-tagged TRIM21 from DEL selection.

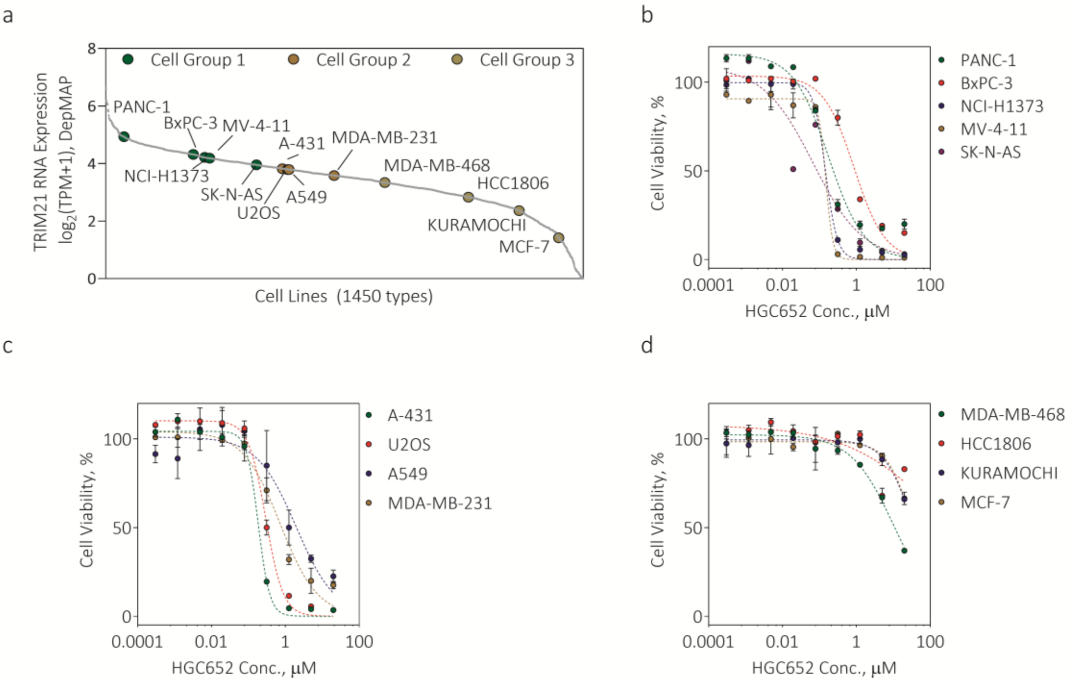


e

| Cell Group 1 | $IC_{50}$, μM | Cell Group 2 | $IC_{50}$, μM | Cell Group 3 | $IC_{50}$, μM |
|---|---|---|---|---|---|
| PANC-1 | 0.146 | U2OS | 0.324 | MDA-MB-468 | 10.95 |
| BxPC-3 | 0.607 | A549 | 0.822 | HCC1806 | >20 |
| NCI-H1373 | 0.135 | A-431 | 0.194 | HURAMOCHI | >20 |
| MV-4-11 | 0.123 | MDA-MB-231 | 0.462 | MCF-7 | >20 |
| SK-N-AS | 0.106 | | | | |

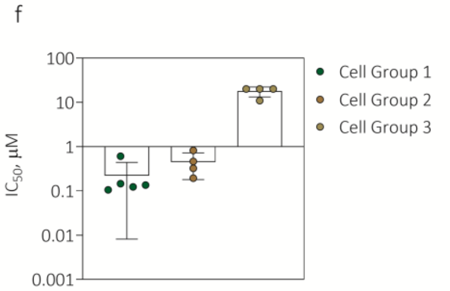


**Figure S2.** Effects of **HGC652** and **HGC645** to multiple cancer cell lines following 72h of compound treatment.

**a,** TRIM21 mRNA expression analysis across selected cancer cell lines. Data of 1450 cancer cell lines were originally retrieved from DepMAP database (https://depmap.org/portal/) and displayed from high expression to low expression. Selected cancer cell lines were grouped into 3 categories representing high expression level (Cell Group 1), medium expression level (Cell Group 2) and high expression level (Cell Group 3). **b,** Growth effects of **HGC652** and **HGC645** to Cell Group 1 including PANC-1, BxPC-3, NCI-H1373, MV-4-11 and SK-N-AS. **c,** Growth effects of **HGC652** and **HGC645** to Cell Group 2 including A-431, U2OS, A549 and MDA-MB-231 cells. **d,** Effects of **HGC652** and **HGC645** to Cell Group 3 including MDA-MB-468, HCC1806, KURAMOCHI and MCF-7 cells. **e,** Derived $IC_{50}$ values of **HGC652** to 13 cancer cell lines. **f,** $IC_{50}$ values of **HGC652** displayed among 3 Cell Group.

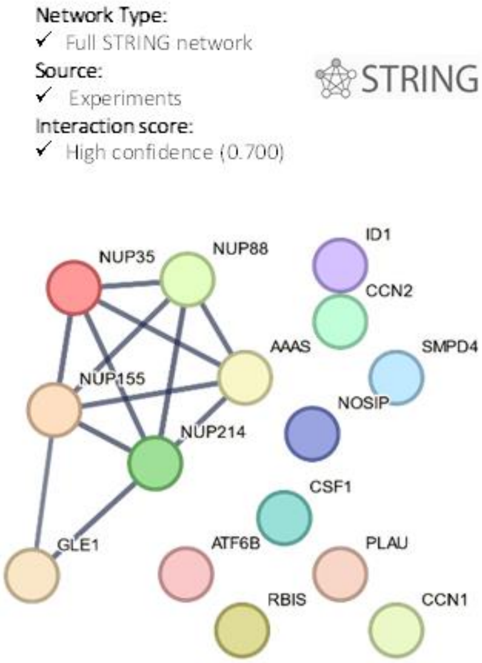


**Figure S3.** Protein-protein interaction networks analysis using the STRING database

### General Procedure for Synthesis of TRIM21-Ligands

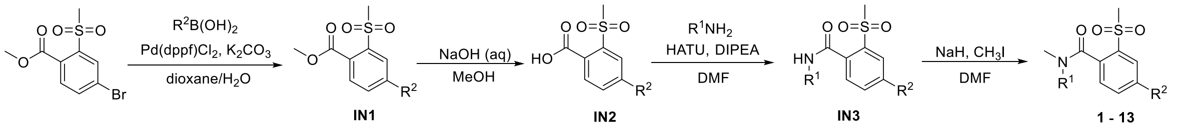


A mixture of methyl 4-bromo-2-(methylsulfonyl)benzoate (1.0 equiv), $R^2$-$B(OH)_2$ (1.2 equiv), $Pd(dppf)Cl_2$ (10 mol%), and $K_2CO_3$ (2 equiv) in 1,4-dioxane/$H_2O$ (4:1, v/v) was stirred under $N_2$ at 100 °C for 16 h. After being cooled to room temperature, the mixture was diluted with water and extracted with EtOAc. The organic layer was washed with brine, dried over anhydrous $Na_2SO_4$, and concentrated in vacuo. The residue was purified by flash chromatography to afford desired product **IN1**.

To a solution of compound **IN1** (1 equiv) in 4:1 mL (v/v) MeOH/$H_2O$ was added NaOH (10 equiv). The resulting mixture was stirred at 60°C for 4 h and the pH was adjusted to 3 by HCl aqueous solution. Then the reaction mixture was diluted with water and EtOAc. The organic layer was separated, washed with brine, dried over anhydrous $Na_2SO_4$, and concentrated in vacuum to afford compound **IN2**. The crude product was used directly in the next step without further purification.

To a solution of compound **IN2** (1 equiv), DIPEA (2 equiv) and HATU (1.2 equiv) in DMF at 0°C was added amine R1-$NH_2$ (1.2 equiv). After 30 min at 0°C, the reaction mixture was diluted with water and EtOAc. The organic layer was separated, washed with water and brine, dried over anhydrous $Na_2SO_4$, and concentrated in vacuum. The crude product was purified by flash chromatography to afford desired product **IN3**.

To a solution of compound **IN3** (1 equiv) in DMF at 0°C was added NaH (1.5 equiv). After 30 min at 0°C, $CH_3I$ (2 equiv) was added to the reaction mixture. The mixture was stirred at room temperature for 2 h and diluted with water and EtOAc. The organic layer was separated, washed with water and brine, dried over anhydrous $Na_2SO_4$, and concentrated in vacuum. The crude product was purified by MPLC to afford desired product **1 – 13**.

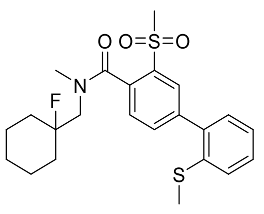

**Compound 1**

**Compound 1**: Light yellow solid, 15.5 mg, 93% yield. Two rotamers were observed by NMR. **$^1$H NMR**

(400 MHz, DMSO-*d*$_6$) δ 7.96 (d, *J* = 2 Hz, 1H), 7.84 – 7.80 (m, 1H), 7.58 (d, *J* = 8.0 Hz, 1H), 7.51-7.38 (m, 2H), 7.36-7.24 (m, 2H), 4.14-3.88 (m, 1H), 3.46-3.34 (m, 1H), 3.29 (s, 3H), 2.88 (s, 3H), 2.42 (s, 3H), 1.86 (br s, 2H), 1.56 (br s, 8H). **LCMS** (ESI$^+$) calculated for $C_{23}H_{28}FNO_3S_2$ m/z [M+H]$^+$: 450.1. Found: 450.0;

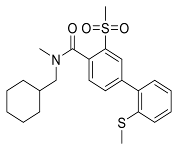

Compound 1a

**Compound 1a**: White solid, 28.9 mg, 43% yield. Two rotamers were observed by NMR. **$^1$H NMR** (400 MHz, DMSO-*d*$_6$) δ 7.94 (d, *J* = 1.6 Hz, 1H), 7.80 (dd, *J* = 8.0, 1.6 Hz, 1H), 7.55-7.39 (m, 3H), 7.35-7.26 (m, 2H), 3.60-3.47, 3.15-3.05 and 2.96-2.82 (m, 2H), 3.29 and 3.26 (s, 3H), 2.98 and 2.79 (s, 3H), 2.42 and 2.41 (s, 3H), 1.85-1.53 (m, 6H), 1.34-1.10 (m, 3H), 1.08-0.59 (m, 2H). **LCMS** (ESI$^+$) calculated for $C_{23}H_{29}NO_3S_2$ m/z [M+H]$^+$: 432.2. Found: 432.1;

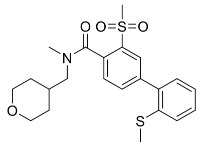

Compound 1b

**Compound 1b**: White solid, 6.0 mg, 39% yield. Two rotamers were observed by NMR. **$^1$H NMR** (400 MHz, DMSO-*d*$_6$) δ 7.94 (d, *J* = 1.6 Hz, 1H), 7.83-7.77 (m, 1H), 7.57-7.51 (m, 1H), 7.50-7.39 (m, 2H), 7.37-7.25 (m, 2H), 3.92-3.73 (m, 2H), 3.55-2.83 (m, 7H), 3.00 and 2.81 (s, 3H), 2.42 and 2.41 (s, 3H), 2.09-1.83 (m, 1H), 1.76-1.42 (m, 2H), 1.34-0.91 (m, 2H). **LCMS** (ESI$^+$) calculated for $C_{22}H_{27}NO_4S_2$ m/z [M+H]$^+$: 434.1. Found: 434.2;

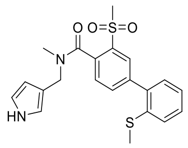

Compound 1c

**Compound 1c**: White solid, 7.1 mg, 36% yield. Two rotamers were observed by NMR. **$^1$H NMR** (600 MHz, DMSO-*d*$_6$) δ 10.70 (d, *J* = 28.8 Hz, 1H), 7.96-7.95 (m, 1H), 7.81-7.79 (m, 1H), 7.63-7.50 (m, 1H), 7.48-7.44 (m, 1H), 7.42-7.40 (m, 1H), 7.34-7.27 (m, 2H), 6.83-6.74 (m, 1H), 6.73-6.71 (m, 1H), 6.11-6.06 (m, 1H), 4.67-4.40 (m, 1H), 4.18-3.02 (m, 1H), 3.33 (s, 3H), 2.88-2.68 (m, 3H), 2.42 (d, *J* = 5.4 Hz, 3H). **LCMS** (ESI$^+$) calculated for $C_{21}H_{22}N_2O_3S_2$ m/z [M+H]$^+$: 415.1. Found: 415.1;

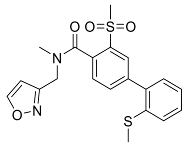

Compound 1d

**Compound 1d**: White solid, 20 mg, 77% yield. Two rotamers were observed by NMR. **$^{1}$H NMR** (400 MHz, DMSO-$d_6$) δ 8.95-8.92 (m, 1H), 7.99-7.94 (m, 1H), 7.85 and 7.78 (dd, *J* = 8.0, 2.0 Hz, 1H), 7.63 and 7.59 (d, *J* = 8.0 Hz, 1H), 7.51-7.39 (m, 2H), 7.36-7.28 (m, 2H), 6.68 and 6.67 (d, *J* = 1.6 Hz, 1H), 4.92-4.34 (m, 2H), 3.34 and 3.33 (s, 3H), 2.97 and 2.81 (s, 3H), 2.42 and 2.41 (s, 3H). **LCMS** (ESI$^+$) calculated for $C_{20}H_{20}N_2O_4S_2$ m/z [M+H]$^+$: 417.1. Found: 417.0;

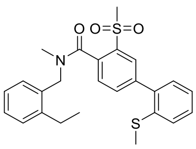

Compound 1e

**Compound 1e**: White solid, 5 mg, 24% yield. Two rotamers were observed by NMR. **$^{1}$H NMR** (400 MHz, DMSO-$d_6$) *δ* 8.01-7.79 (m, 2H), 7.68-7.58 (m, 1H), 7.50-7.36 (m, 3H), 7.35-7.13 (m, 5H), 5.03, 4.48 (d, *J* = 15.2 Hz) and 4.37 (s, 2H), 3.34 (s, 3H), 2.98 and 2.72 (s, 3H), 2.74-2.65 and 2.45-2.32 (m, 5H), 1.21 and 0.93 (t, *J* = 7.6 Hz, 3H). **LCMS** (ESI$^+$) calculated for $C_{25}H_{27}NO_3S_2$ m/z [M+H]$^+$: 454.1. Found: 454.2;

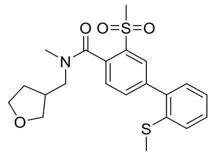

Compound 1f

**Compound 1f**: White solid, 7.1 mg, 25% yield. Two rotamers were observed by NMR. **$^{1}$H NMR** (400 MHz, DMSO-$d_6$) δ 7.95 (d, *J* = 2.0 Hz, 1H), 7.81 (dd, *J* = 8.0, 2.0 Hz, 1H), 7.63-7.53 (m, 1H), 7.50-7.40 (m, 2H), 7.36-7.26 (m, 2H), 3.88-3.76 and 3.14-3.05 (m, 2H), 3.73-3.63 (m, 1H), 3.61-3.42 (m, 3H), 3.32-3.26 (m, 3H), 3.04-2.79 (m, 3H), 2.73-2.55 (m, 1H), 2.46-2.39 (m, 3H), 2.09-1.84 (m, 1H), 1.78-1.44 (m, 1H). **LCMS** (ESI$^+$) calculated for $C_{21}H_{25}NO_4S_2$ m/z [M+H]$^+$: 420.1. Found: 420.0;

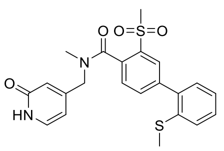

Compound 1g

**Compound 1g**: Yellow solid, 2.7 mg, 11% yield. Two rotamers were observed by NMR. **$^{1}$H NMR** (400

MHz, DMSO-$d_6$) δ 11.50 (s, 1H), 7.98 and 7.97 (d, $J$ = 2.0 Hz, 1H), 7.85 and 7.78 (dd, $J$ = 8.0, 2.0 Hz, 1H), 7.65 and 7.57 (d, $J$ = 8.0 Hz, 1H), 7.51-7.39 (m, 2H), 7.39-7.26 (m, 3H), 6.40-6.13 (m, 2H), 4.55 and 4.16 (m, 2H), 3.35-3.30 (m, 3H), 2.94 and 2.78 (s, 3H), 2.43 and 2.41 (s, 3H). **LCMS** calculated for $C_{22}H_{22}N_2O_4S_2$ m/z $[M+H]^+$: 443.1. Found: 443.1;

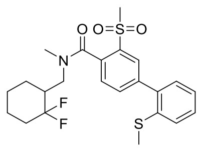

Compound 2a

**Compound 2a**: White solid, 7.4 mg, 55% yield. Two rotamers were observed by NMR. **$^1$H NMR** (400 MHz, DMSO-$d_6$) δ 7.95 (d, $J$ = 2.0 Hz, 1H), 7.82 (dd, $J$ = 8.0, 2.0 Hz, 1H), 7.62-7.39 (m, 3H), 7.37-7.26 (m, 2H), 3.90-3.47 (m, 1H), 3.31-3.24 (m, 4H), 3.03-2.77 (m, 3H), 2.44-2.38 (m, 3H), 2.14-1.59 (m, 5H), 1.53-0.92 (m, 4H). **LCMS** ($ESI^+$) calculated for $C_{23}H_{27}F_2NO_3S_2$ m/z $[M+H]^+$: 468.1. Found: 468.0;

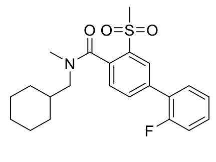

Compound 2b

**Compound 2b**: White solid, 17 mg, 65% yield. Two rotamers were observed by NMR. **$^1$H NMR** (400 MHz, DMSO-$d_6$) δ 8.10 (s, 1H), 8.02-7.95 (m, 1H), 7.74-7.64 (m, 1H), 7.59-7.48 (m, 2H), 7.44-7.35 (m, 2H), 3.62-3.45 3.20-3.00 and 2.96-2.82 (m, 2H), 3.31 and 3.28 (s, 3H), 2.98 and 2.79 (s, 3H), 1.91-1.69 (m, 4H), 1.67-1.50 (m, 2H), 1.29-1.11 (m, 3H), 1.10-0.63 (m, 2H). **LCMS** ($ESI^+$) calculated for $C_{22}H_{26}FNO_3S$ m/z $[M+H]^+$: 404.2. Found: 404.2;

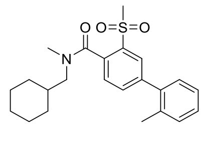

Compound 2c

**Compound 2c**: White solid, 8.1 mg, 26% yield. Two rotamers were observed by NMR. **$^1$H NMR** (400 MHz, DMSO-$d_6$) δ 7.88-7.83 (m, 1H), 7.83-7.75 (m, 1H), 7.55-7.47 (m, 1H), 7.39-7.26 (m, 4H), 3.61-3.48 3.16-3.04 and 2.94-2.84 (m, 2H), 3.32 and 3.28 (s, 3H), 2.98 and 2.80 (s, 3H), 2.30-2.23 (m, 3H), 1.89-1.47 (m, 6H), 1.35-1.10 (m, 3H), 1.09-0.61 (m, 2H). **LCMS** ($ESI^+$) calculated for $C_{23}H_{29}NO_3S$ m/z $[M+H]^+$: 400.2. Found: 400.1;

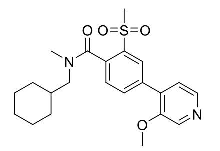

Compound 2d

**Compound 2d:** White solid, 9.3 mg, 28% yield. Two rotamers were observed by NMR. **$^1$H NMR** (400 MHz, DMSO-$d_6$) δ 8.92-8.21 (m, 2H), 8.14 (d, *J* = 2.0 Hz, 1H), 8.04-7.92 (m, 1H), 7.62-7.39 (m, 2H), 3.95 (s, 3H), 3.58-3.47 3.14-3.04 and 2.95-2.81 (m, 2H), 3.30 and 3.27 (s, 3H) 2.97 and 2.79 (s, 3H), 1.91-1.48 (m, 6H), 1.31-1.10(m , 2H), 1.10-0.87 (m, 2H), 0.77-0.61 (m, 1H). **LCMS** (ESI$^+$) calculated for $C_{22}H_{28}N_2O_4S$ m/z [M+H]$^+$: 417.2. Found: 417.2;

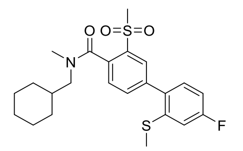

Compound 2e

**Compound 2e**: White solid, 12.4 mg, 71% yield. Two rotamers were observed by NMR. **$^1$H NMR** (600 MHz, DMSO-*$d_6$*) δ 7.92 (d, *J* = 1.8 Hz, 1H), 7.79 (dd, *J* = 7.8, 1.8 Hz, 1H), 7.53 and 7.52 (d, *J* = 7.8 Hz, 1H), 7.41-7.34 (m, 1H), 7.24 (dd, *J* = 10.2, 2.4 Hz, 1H), 7.12 (td, *J* = 8.4, 2.4 Hz, 1H), 3.59-3.49 3.13-3.04 and 2.95-2.82 (m, 2H), 3.30 and 3.26 (s, 3H), 2.98 and 2.79 (s, 3H), 2.47 and 2.46 (s, 3H), 1.87-1.52 (m, 6H), 1.30-1.13 (m, 3H), 1.10-0.64 (m, 2H). **$^{13}$C NMR** (150 MHz, DMSO-*$d_6$*) δ 168.8, 163.9, 162.1, 140.4, 140.2, 137.2, 136.8, 135.1, 134.9, 133.9, 132.1, 130.2, 128.6, 128.2, 112.4, 112.0, 57.5, 53.1, 45.3, 38.2, 36.0, 35.8, 33.0, 30.9, 26.4, 25.8, 15.4. **LCMS** (ESI$^+$) calculated for $C_{23}H_{28}FNO_3S_2$ m/z [M+H]$^+$: 450.2. Found: 450.5;

**NMR Spectrum for Compound 2e**

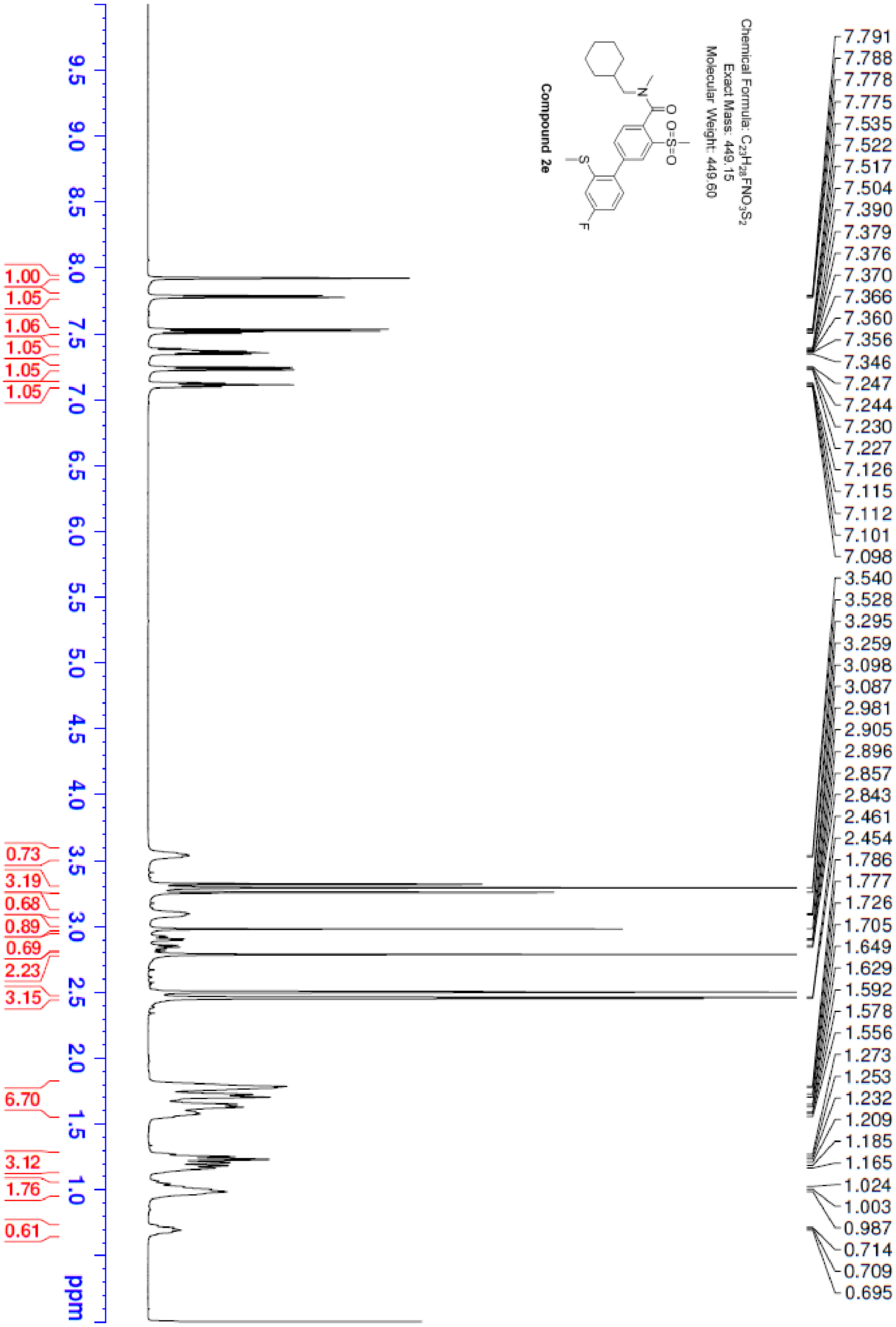

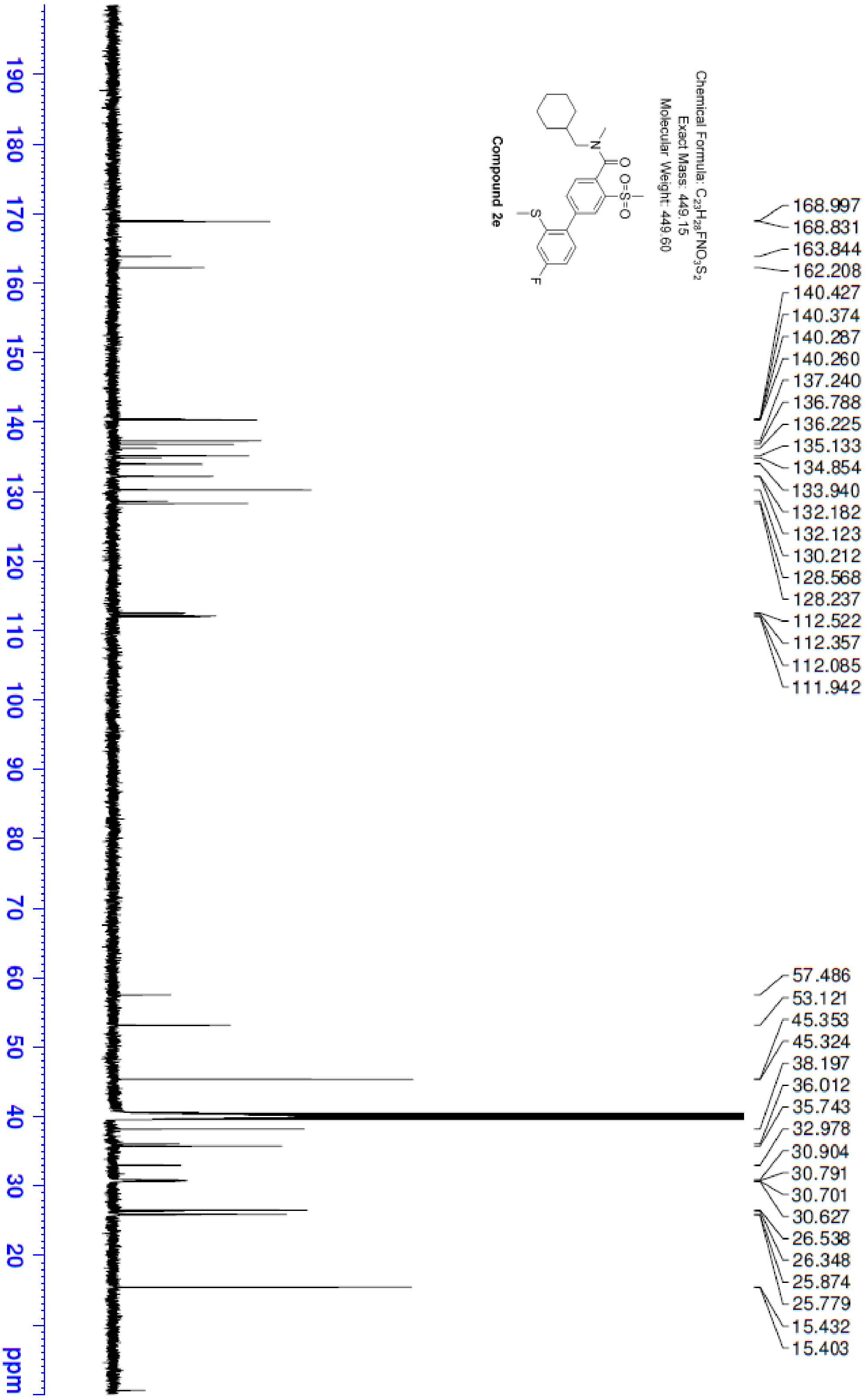
Chemical Formula: $C_{23}H_{28}FNO_3S_2$
Exact Mass: 449.15
Molecular Weight: 449.60
Compound 2e
168.997
168.831
163.844
162.208
140.427
140.374
140.287
140.260
137.240
136.788
136.225
135.133
134.854
133.940
132.182
132.123
130.212
128.568
128.237
112.522
112.357
112.085
111.942
57.486
53.121
45.353
45.324
38.197
36.012
35.743
32.978
30.904
30.791
30.701
30.627
26.538
26.348
25.874
25.779
15.432
15.403
190
180
170
160
150
140
130
120
110
100
90
80
70
60
50
40
30
20
ppm

**LC-MS Spectrum for Compound 2e**

<Chromatogram>

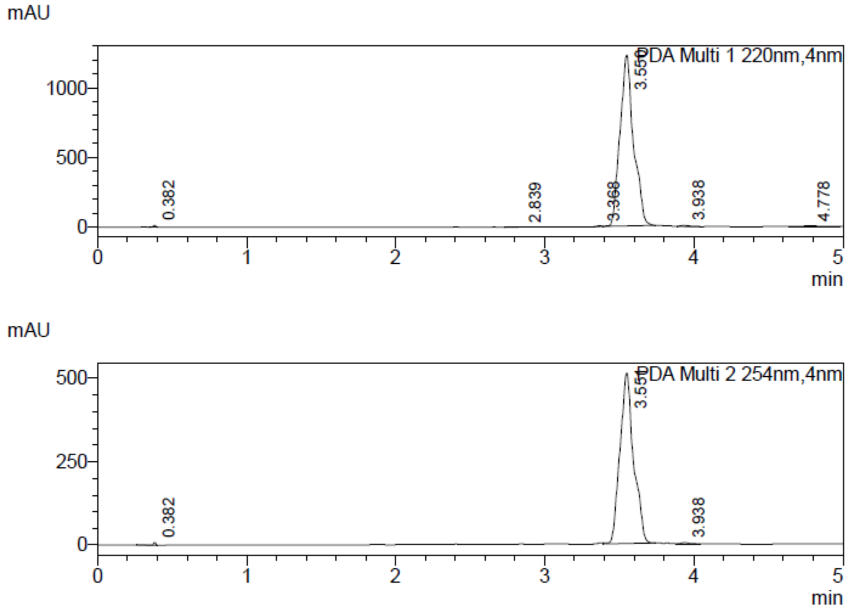


min

<MS Chromatogram>

0.03min delay

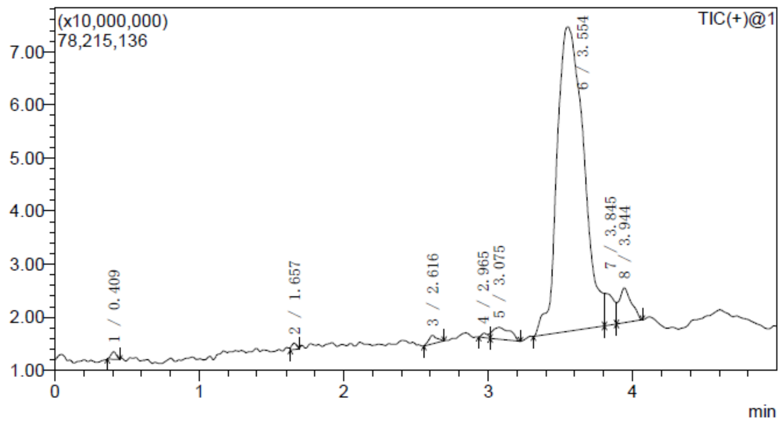


Line#:6
Spectrum Mode:(Averaged==Averaged)3.550-3.560

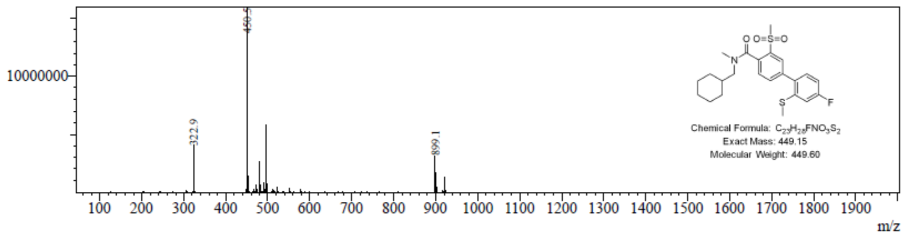

**HPLC Spectrum for Compound 2e**

<Chromatogram>

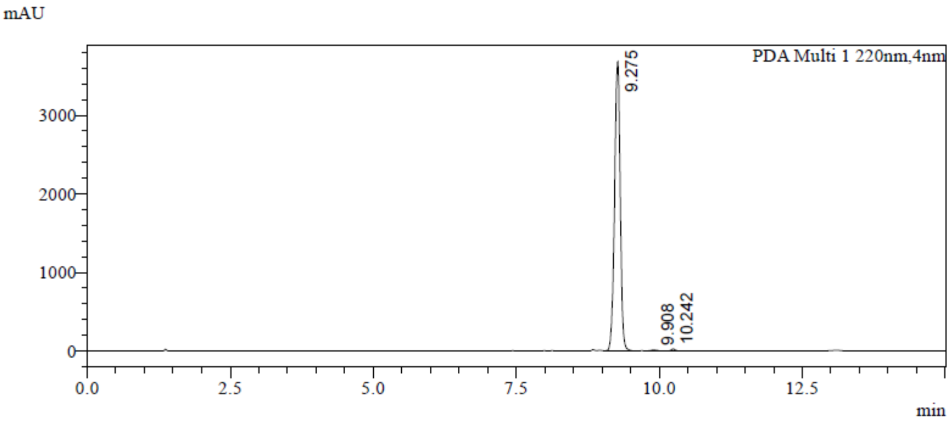


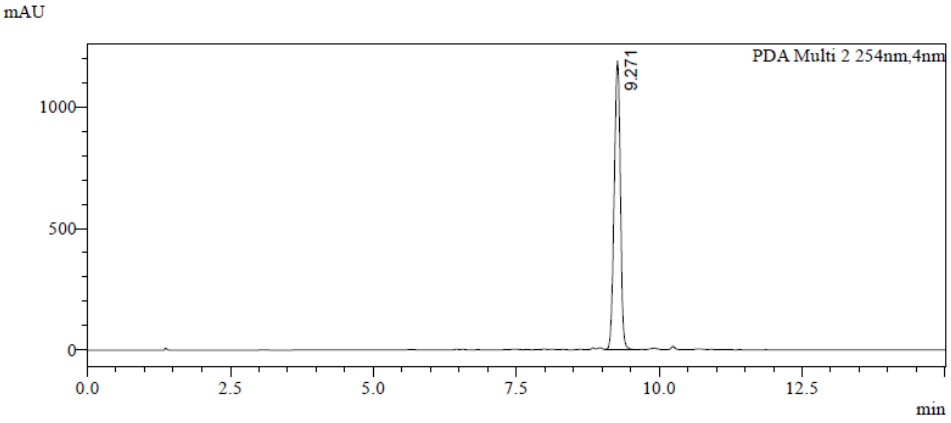


<Chromatogram Peak Table>

Peak Table

PDA Ch1 220nm

| Peak | Ret. Time | Area | Height | Talling.Factor | Resolution | Area% |
|---|---|---|---|---|---|---|
| 1 | 9.275 | 25503874 | 3689973 | 0.920 | -- | 99.076 |
| 2 | 9.908 | 111648 | 12570 | -- | 2.838 | 0.434 |
| 3 | 10.242 | 126184 | 26744 | 1.138 | 1.696 | 0.490 |
| 总计 | | 25741705 | 3729287 | | | 100.000 |

PDA Ch2 254nm

| Peak | Ret. Time | Area | Height | Talling.Factor | Resolution | Area% |
|---|---|---|---|---|---|---|
| 1 | 9.271 | 8911315 | 1187858 | 0.947 | -- | 100.000 |
| 总计 | | 8911315 | 1187858 | | | 100.000 |
